# Molecular free energy optimization on a computational graph

**DOI:** 10.1101/2020.04.01.020214

**Authors:** Xiaoyong Cao, Pu Tian

## Abstract

Free energy is arguably the most important property of molecular systems. Despite great progress in both its efficient estimation by scoring functions/potentials and more rigorous computation based on extensive sampling, we remain far from accurately predicting and manipulating biomolecular structures and their interactions. There are fundamental limitations, including accuracy of interaction description and difficulty of sampling in high dimensional space, to be tackled. Computational graph underlies major artificial intelligence platforms and is proven to facilitate training, optimization and learning. Combining autodifferentiation, coordinates transformation and generalized solvation free energy theory, we construct a computational graph infrastructure to realize seamless integration of fully trainable local free energy landscape with end to end differentiable iterative free energy optimization. This new framework greatly improves efficiency by replacing local sampling with differentiation. Its specific implementation in protein structure refinement achieves superb efficiency and competitive accuracy when compared with state of the art all-atom mainstream methods.

## Introduction

Full understanding of free energy landscape (FEL) for a given molecular system implicates the ability to accurately predicting its behavior, and provides a rational basis for further manipulation and design. Scientists have made tremendous effort in calculation of FEL with great achievements in advancement of both theory and computational algorithms.^1^ Rigorous computation of FEL usually involves sampling of configurational space by molecular simulations and post-processing of generated trajectories/statistics. More efficient estimation as utilized in design/prediction/refinement of protein structures^2–4^ involves repetitive proposal/sampling and/or energy minimization of candidate structures/sequences (e.g. FastRelax^5^) followed by evaluating/scoring with various forms of potentials.^6–8^ All these schemes have fundamental limitations as briefed below:

1. Pairwise approximation of non-bonded molecular interactions is utilized in essentially all molecular modeling with either physics or knowledge based force fields (FF) (e.g. CHARMM,^9^ Rosetta^10^). Two non-bonded basic units (atoms or coarse grained particles) are assumed to have interactions determined only by the distance (and orientation in case of anisotropic units) between them, regardless of identity and spatial distribution of other neighboring units.
2. Fixed simple functional form (e.g. Lennard-Jones, quadratic form) are utilized in traditional FF for convenience of fitting. While having well grounded physical underpinning when relevant basic units are near equilibrium positions (local minima), these functions create a ceiling of accuracy for description of frustrated molecular systems^11^ with a significant fraction of comprising units off energy minima position and/or orientation.
3. Repetitive local sampling is universal. The number of energetically favorable local configurations of a given composition for typical molecular systems (e.g. water, protein) at specific conditions (e.g. temperature, pressure) is a tractable number and form a local FEL (LFEL). The reason is that local correlations are strong in soft condensed matter, thus effective local dimensionality is much smaller than that corresponds to the number of degrees of freedom (DOFs). Different local compositions, which is also a tractable number for similar reason, form different LFEL. Competition and superposition of these overlapping LFEL constitutes global FEL of a given molecular system. However, sampling of these local arrangements are repeatedly carried out with tremendous wasting of computational resources.
4. No direct manipulation of molecular coordinates based on free energy is available for sampling based methods.
5. Maintenance of rigid constraints (e.g. bond lengths) is frequently utilized to improve efficiency of molecular simulations with either shake,^12^ rattle^13^ or settle^14^ algorithm. These iterative procedures maintain bond lengths (and angles) within a preset tolerance off target value, engender computational cost, and may diverge when large forces are experienced by relevant units. Specialized methods, such as concerted rotation^15^ and backrub,^16^ are helpful in maintaining constraints for stochastic configurational space sampling of (bio)polymers. However, these procedures may not be directly driven by differentiation w.r.t. (with respect to) a given potential.

Among these limitations, the first two have fundamental impact on accurate interaction description, the third and the fourth severely decrease computational efficiency, and the fifth is a nuisance. Neural network based many body potentials^17^ may help with the first two limitations. The remaining three are yet to be overcome. We previously developed generalized solvation free energy (GSFE) theory.^8^ The local maximum likelihood approximation (LMLA) of GSFE theory was implemented with a simple neural network to assess protein structural models, and demonstrated strong competitiveness when compared with state-of-the-art knowledge based potentials. The physical essence of GSFE is caching of LFEL by training with data sets derived from high resolution experimental structures. The connection of LFEL to mainstream enhanced sampling and coarse-graining algorithms are described in detail elsewhere.^18^

A graph is a set of vertices and edges connecting them. In a computational graph, vertices are functions that processing data transmitted through edges. Autodifferentiation (AD) is a powerful way of solving exact derivatives first developed in 1974,^19^ its special form of backpropatation (BP) in neural network was reinvented by Hinton ^20^ in 1986. Derivatives, as long as they exists, in all programmable computational process may be calculated with autodifferentiation without explicit functional forms. The gorgeous capability of computational graph empowered by autodifferentiation has been proven by widespread application of major artificial intelligence platforms (e.g. TensorFlow and PyTorch). Integrating the GSFE theory, coordinates transformation and autodifferentiation, we map molecular free energy optimization onto a computational graph to address the last three of above mentioned challenges. Implementation of this algorithm in protein structure refinement (PSR) is demonstrated to be competitive in accuracy and orders of magnitude more efficient when compared with present mainstream methodologies.

## Results

### Mapping of free energy optimization onto a computational graph

With a simple probabilistic description of molecular systems, GSFE formulates a directed link from molecular coordinates to LFEL, superposition of which under strict implicit global correlation restraints (see Equation 3 and Fig.1) is utilized to approximate total free energy of interested molecular system. Caching of LFEL through training of the neural network is described perviously^8^ (and briefed in *Methods* section as well). However, GSFE does not provide measures to efficiently update structures. In principle, with establishment of a computational graph, BP operation of AD from calculated approximate free energy (AFE) to coordinates may provide gradients (and higher ordered derivatives when needed) of AFE w.r.t. coordinates, which may subsequently be updated by a simple gradient descent (or higher ordered) optimization. AD is implemented by major AI platforms and may be turned on by a simple statement. One concern is that in such direct update of coordinates, bond lengths/angles would be changed and one has to utilize iterative algorithms such as shake^12^ or rattle^13^ to maintain constraints if desire. To overcome this issue, we design a coordinate transformation procedure that updates internal coordinates where only derivatives w.r.t. interested quantity (dihedrals) are utilized and derivatives w.r.t. constraint quantity (bond lengths and angles) are set to zero. This way, constraints are realized exactly with no concern of divergence. As shown in Fig. 2, to facilitate feature extraction, a transformation from internal to cartesian coordinates is necessary.

**Figure 1:**
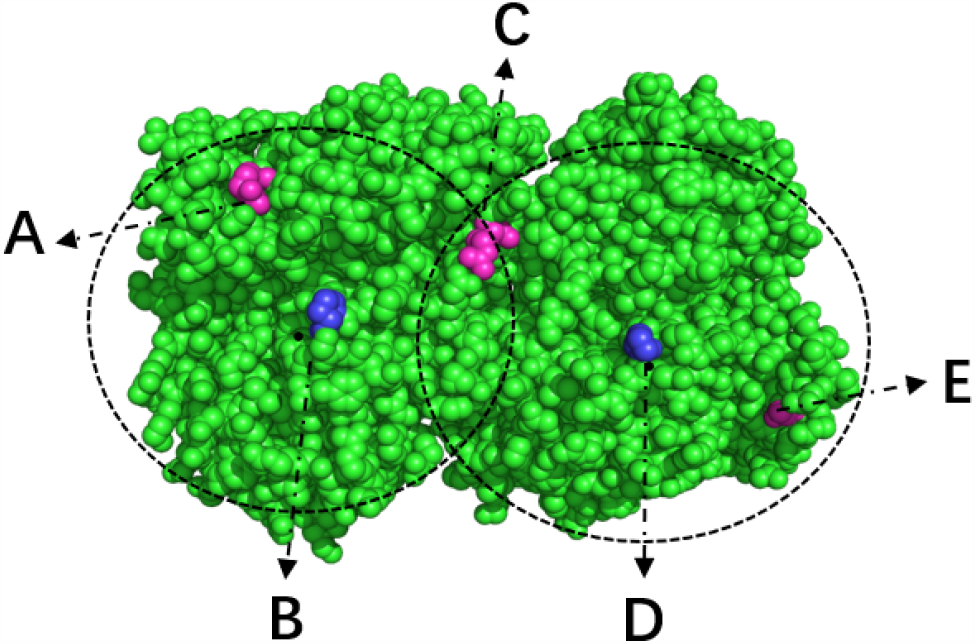
Illustration of implicit mediated global correlations and effective larger cutoff in GSFE-Refinement. Two dashed circles are two “local” region for LFEL centered at solute unit B and D respectively. As a comprising unit of LFELs centered on both unit B and D, unit C experience effective force from unit A as mediated by B, and effective force from E as mediated by D. In fact, each unit experience effective forces mediated by LFELs defined by all of its solvent unit, resulting in an effective large cutoff that is approximately two times the radius of dashed circle for defining LFEL as shown. All mediated global correlations in essence is the equality of shared states for overlapping DOFs belong to different LFELs, since only one set of coordinates is used in the whole optimization process, these correlations are naturally maintained.

**Figure 2:**
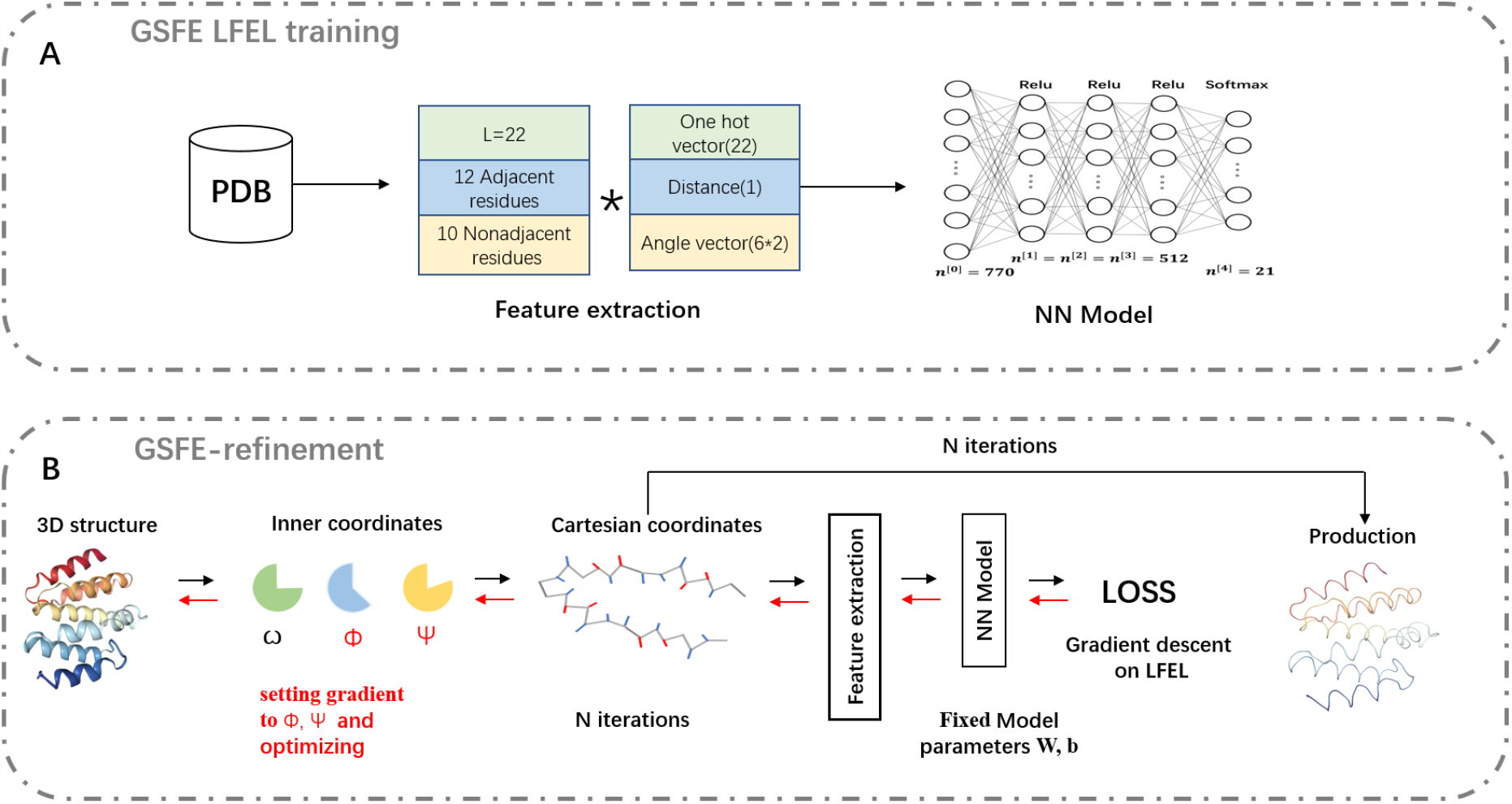
Schematic representation of GSFE-Refinement training and optimization. (A)illustration of LFEL training based on LMLA-GSFE(see Ref ?? for details). (B) Illustration of the fully end-to-end differentiable optimization pipeline. Small solid black arrows represent forward pass computation of approximate free energy, Small solid red arrows represent BP operation for taking derivatives. The empty arrow represents input of decoys. Final result is delivered as output after a preset iteration number N.

### GSFE-Refinement performance on 3DRobot data set

#### Refinement with LMLA-GSFE

In order to investigate the efficiency of optimization with BP operation of AD and the accuracy of LMLA-GSFE, we carried out refinement with 770LFEL on the 3DRobot dataset (see *Methods*). The results for the best of top 5 models at learning rates of 0.001,0.0005 and 0.0001 are evaluated by *C*_*α*_ RMSD(root-mean-squared deviation) and GDT-HA (global distance test high accuracy) as indicators^21^ and listed in Table 1(rows 1,2,3). Avg-ΔGDT-HA (see Table 1) with learning rates 0.001,0.0005 and 0.0001 are −1.38,-0.18 and 0.2, corresponding GDT-HA-num (see Table 1) are 95/322,134/322 and 182/322 respectively. Within limited range of examination, we observe on average the smaller the learning rate, the larger the average ΔGDT-HA and the larger the GDT-HA-num (as shown in Fig. 3A and Fig. 4A,B,C). However, for some decoys, a larger learning rate improves refinement (see Fig.S8). Physically, a larger learning rate means refinement with a larger step in all of considered LFEL and costs less computing resource. Considering all these factors, a learning rate of 0.0005 is utilized hereafter.

**Table 1:** Summary for the best of top 5 models with various combinations for LR/*λ*/W on 3DRobot dataset

| Row | LR/$\lambda$/W<sup>a</sup> | Avg-ΔGDT-HA<sup>b</sup> | GDT-HA-num<sup>c</sup> | Avg-ΔRMSD<sup>d</sup> | RMSD-num<sup>e</sup> |
| --- | --- | --- | --- | --- | --- |
| 1 | 0.001/0/0 | -1.38 | 95/322 | 0.0022 | 130/322 |
| 2 | 0.0005/0/0 | -0.18 | 134/322 | -0.0175 | 158/322 |
| 3 | 0.0001/0/0 | 0.2 | 182/322 | -0.0099 | 201/322 |
| 4 | 0.0005/0.1/0 | -0.19 | 137/322 | -0.0168 | 166/322 |
| 5 | 0.0005/1.2/0 | -0.02 | 162/322 | -0.0147 | 188/322 |
| 6 | 0.0005/10.0/0 | 0.24 | 211/322 | -0.0167 | 257/322 |
| 7 | 0.0005/0.1/1 | 0.02 | 163/322 | -0.0136 | 191/322 |
| 8 | 0.0005/1.2/1 | 0.26 | 214/322 | -0.0178 | 275/322 |
| 9 | 0.0005/10.0/1 | 0.27 | 210/322 | -0.019 | 291/322 |

<sup>a</sup>LR is the learning rate, $\lambda$ is the coefficient of smooth<sub>l1</sub> loss for conformation restraints, W is the AA weight (see Equations 5 and 6, with 1 represents on and 0 represents off.)
<sup>b</sup>The average value of ΔGDT-HA for all decoys
<sup>c</sup>The average number of decoys with ΔGDT-HA>0 for all 36 decoy sets
<sup>d</sup>The average value of ΔRMSD for all decoys
<sup>e</sup>The average number of decoys with ΔRMSD<0 for all 36 decoy sets

**Figure 3:**
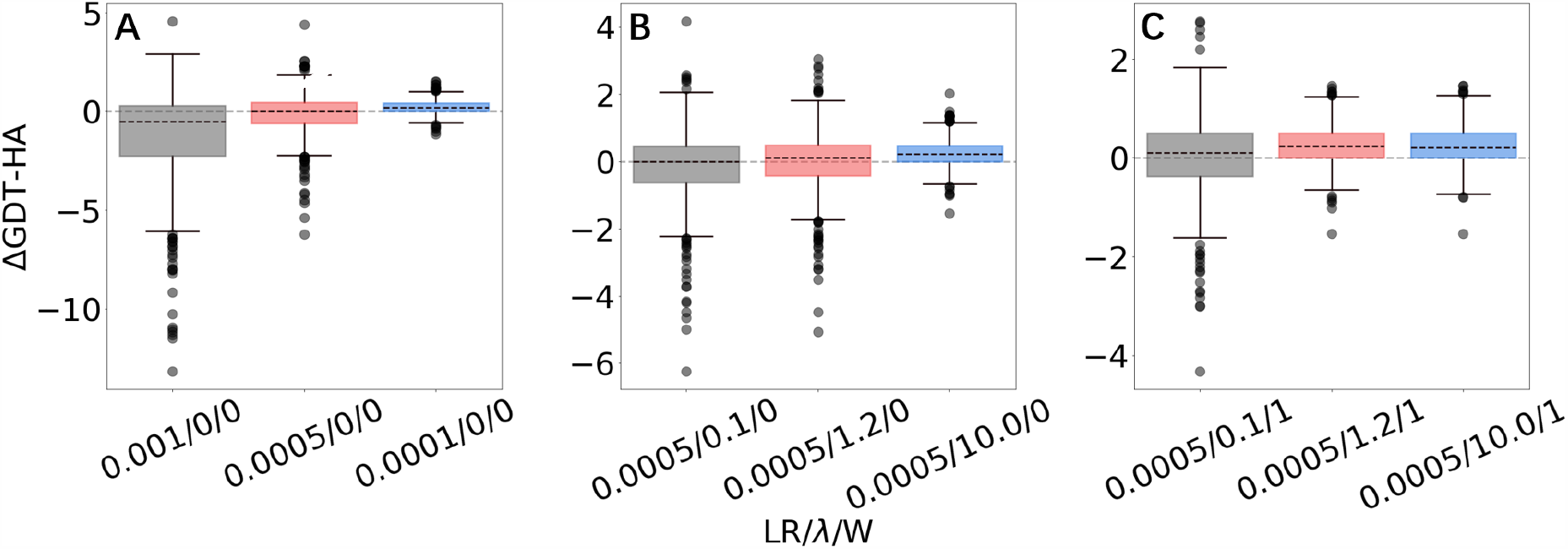
Box plots for ΔGDT-HA with different LR/*λ*/W combinations for 3DRobot dataset. More box plots of ΔRMSD and ΔGDT-HA are available in supporting information (Fig. S1,2,3).

**Figure 4:**
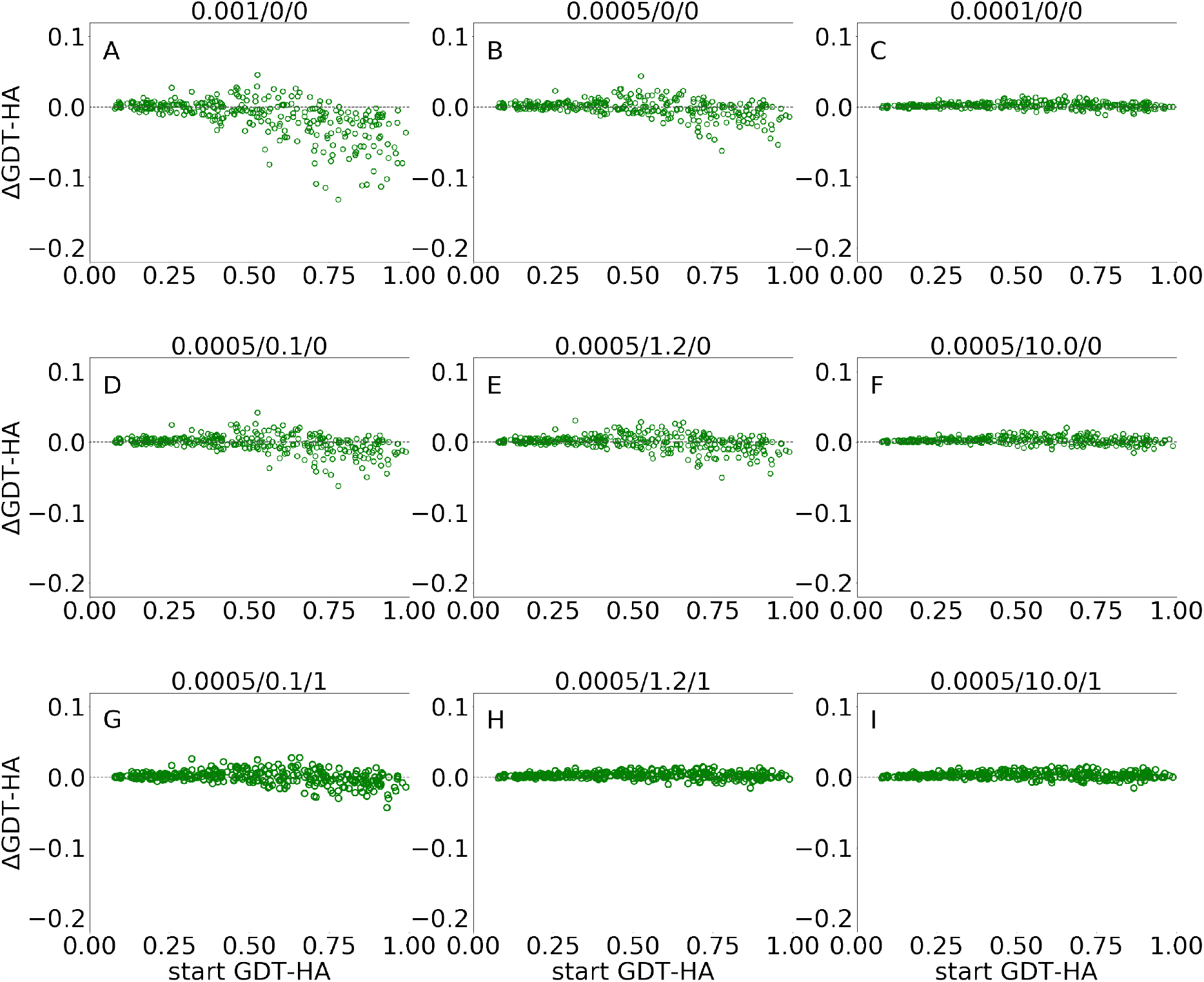
Scatter plots of ΔGDT-HA as a function of start GDT-HA score for best of top5 models from GSFE-Refinement for 3DRobot dataset. Corresponding LR/*λ*/W combination is noted on top of each plot. More scattered plots of ΔRMSD and ΔGDT-HA are available in supporting information (Fig. S4,5,6).

#### Impact of the Smooth_l1 term

In view of the increasing accuracy of the starting structures generated by protein structure prediction, the imposition of reasonable restrictions has become an important part of refinement.^22–25^ Here, we add the smooth_l1 loss term (see Equations 4 and 7 in *Methods*) to limit the conformation search space to be in the vicinity of starting structures. As shown in Table 1 (rows 4,5 and 6), the Avg-ΔGDT-HA for *λ* = 0.1, 1.2, 10.0 are −0.19, −0.02, 0.24 and corresponding GDT-HA-num are 137/322,162/322 and 211/322 respectively. The Avg-ΔGDT-HA based on *λ* = 10.0 is significantly better than that based on *λ* = 0.1 and 1.2, and GDT-HA-num and RMSD indicators show similar trends (as shown in Fig. 3.B and Fig. 4D,E,F).

#### Impact of local weights on the refinement performance

There have been some studies on local restraints in protein structure refinement based on prior knowledge,^22,26,27^ selection of specific regions,^22,26,28^ and local structure evaluation.^22,26,29^ In our training of LFEL, the available data are significantly different for each of amino acids (AA). Larger datasets are highly likely to improve description of LEFL surrounding corresponding AAs. To test this speculation, the fraction of each AA is used as the refinement weight for LFEL (see Equations 5, 6). Table 1 shows the significantly improved results after adding local weights (rows 7, 8 and 9 compared with rows 4, 5 and 6). For example, when LR/*λ*/W changes from 0.0005/1.2/0 to 0.0005/1.2/1, the Avg-ΔGDT-HA increased from −0.02 to 0.26 (row 5 and row 8). Various other evaluation indicators (GDT-HA-num, RMSD and RMSD-num) exhibit similar improvement (as shown in Fig. 3 B,C and Fig. 4D,E,F,G,H,I).

### GSFE-Refinement performance on refineD data set

In order to further investigate the robustness of GSFE-Refinement, we test its performance on the refineD dataset at LR/*λ*/W = 0.0005/1.2/1 with 770-LFEL. As shown in Table 2 (see detailed results in Table S1 and S2 in support information), in top 1 models, the GDT-HA score of GSFE-Refinement is 0.08 and ranks the fourth. In best of top5 models, GSFE-Refinement ranks the sixth, its result (0.4400) is better than FastRelax-0.5Å(0.0548), FastRelax-4.0Å(0.0751), FastRelax (−0.1999), ModRefiner-0 (−0.8400), and ModRefiner-100 (0.1491). These results are generated by 5 iterations within a few seconds on a single CPU core, in strong contrast to conventional sampling and minimization (e.g. FastRelax^5^) where thousands or even hundreds of thousands of iterations and hours of CPU time are usually necessary. It is also important to note that despite all heavy atom in side chains other than *C*_*β*_ are missing in our model, competitive accuracy is achieved when compared with state-of-the-art full heavy atom methods. However, the superior efficiency is mainly due to substitution of local sampling by differentiation. The number of atoms in GSFE-Refinement is more than half of all heavy atoms.

**Table 2:** Summary of GSFE-Refinement and other refinement methods on the 150-target refineD dataset (results for other methods are taken from Ref^30^)

| Method | Avg.Top 1 | Avg.Best of 5 | GDT-HA-num |  |
| --- | --- | --- | --- | --- |
| refined-C | 0.6365 | 1.3109 | 121/150 |  |
| refined-NC | -1.2403 | 1.5343 | 104/150 |  |
| FG-MD | 0.5597 | 0.5597 | - |  |
| FastRelax | -3.4317 | -0.1999 | - |  |
| FastRelax-0.5Å | -0.3411 | 0.8811 | 90/150 | More details for top1 and the |
| FastRelax-2.0Å | -1.2120 | 0.8223 | 77/150 |  |
| FastRelax-4.0Å | -2.5471 | 0.0751 | 67/150 |  |
| ModRefiner-0 | -0.8400 | -0.8400 | - |  |
| ModRefiner-100 | 0.1491 | 0.1491 | - |  |
| GSFE-Refinement | 0.0800 | 0.4400 | 112/150 |  |

best of top 5 models for refinedD data set are available in supporting information (Table S1,2).

### GSFE-Refinement performance on CASP 11 and CASP12 data set

GSFE-Refinement is further evaluated on the CASP11 and CASP12 dataset (see *Methods*). Based on above mentioned results, we utilize a parameter combination LR/*λ*/W = 0.0005/1.2/1. In Fig. 5, we present structural change of the CAPS11 and CASP12 decoys after GSFE-Refinement as measured by ΔGDT-HA and ΔRMSD, and improvement is observed for most of the cases. Specifically, there are 50%(17/34) and 88.2%(30/34) successful refinement when evaluated by GDT-HA and RMSD scores in CASP11 (as shown in Fig. 5A, B), and 64.5%(20/31) and 100%(31/31) when evaluated by GDT-HA and RMSD scores in CASP12 (as shown in Fig. 5C,D). (see detailed results in Table S3,S4,S5, S6 and Fig. S7 in support information).

**Figure 5:**
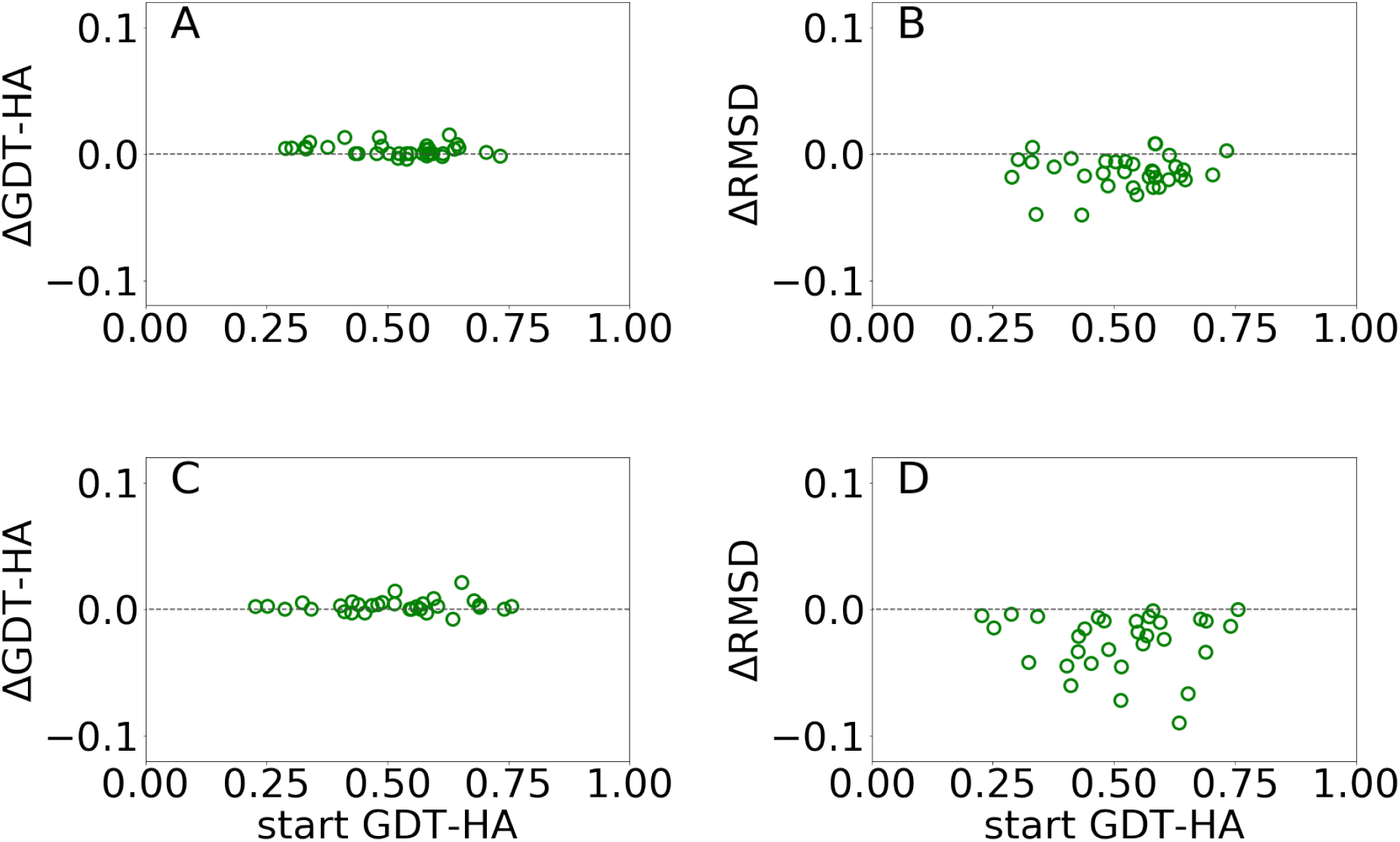
ΔGDT-HA and ΔRMSD of best of top 5 models as a function of start GDT-HA score obtained from GSFE-Refinement of CASP11 (A) and (B) and CASP12 (C) and (D) datasets. Corresponding plots for top 1 models are presented in supporting information (Fig. S7).

### GSFE-Refinement assessment in CASP 14

We participated in CASP14 competition and submitted 180 models for 36 targets (we registered late and missed the first 13 targets), among which 31 are effective targets in CASP14 final statistics. As shown in Fig.6, we improved 38.7% targets compared with the 24.8% of CASP14 average as measured by ΔGDT-HA, and improved 38.7% targets compared with the 27.7% average as measured by ΔRMSD_CA. In top 1 model, GSFE-Refinement(with GR code 294) ranks 12 according to SUM Zscore(>0.0) based on GDT-TS score. And AVG Zscore(>0.0) based on GDT-TS score ranks 6 (https://predictioncenter.org/casp14/zscores_final_refine.cgi). Specific Z scores of GSFE-Refinement is provided in supporting Fig. S9. In particular, for 13 targets with start GDT-TS score better than 60, GSFE-Refinement ranks the first. Again, we performed 5 iterations for each target, with computing cost ranging from 0.7 to 2.7 seconds on a single CPU core.(R1028 has 75 amino acids and costs 0.7s. R1042v1 contains 276 amino acids, which takes 2.7s)

**Figure 6:**
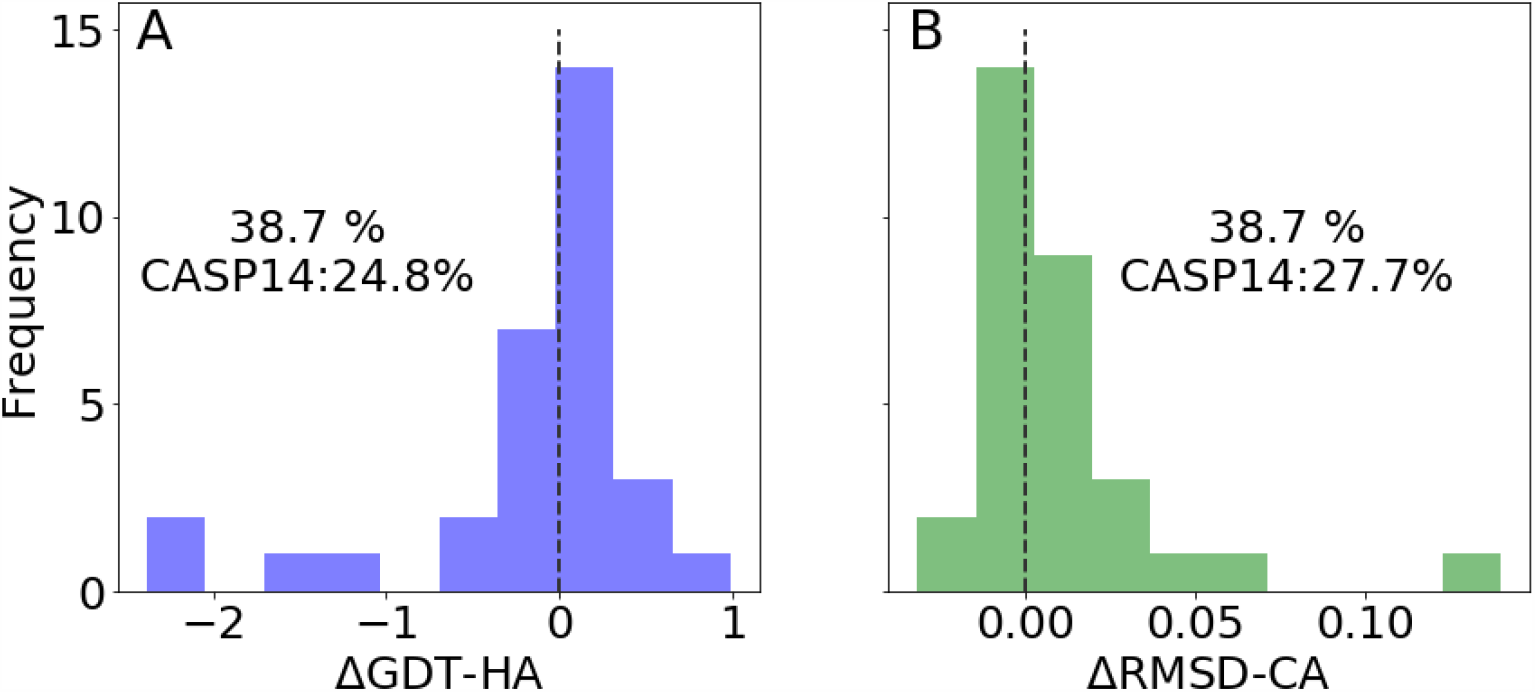
GSFE-Refinement performance in CASP14. Success (percentage of improved targets for selected indicators) rate of GSFE-Refinement and CASP14 average are shown. (A) Distribution of ΔGDT-HA. (B) Distribution of ΔRMSD.

## Discussions

Optimization of molecular free energy on a computational graph proposed and demonstrated in this work makes two distinct contributions,

1. Replacement of expensive local sampling by differentiation w.r.t. cached LFEL realizes superb efficiency.
2. Combination of AD and coordinate transformation realizes exact hard constraints with minimal computational cost.

GSFE-Refinement takes a few seconds on a single CPU core for refinement of typical decoys, in strong contrast to hours for typical sampling and energy minimization with knowledge based potential (e.g. FastRelax^5^) without explicit water representation, and thousands or even tens of thousands hours for refinement based on MD simulations with explicit water representation. As backbone and *C*_*β*_ atoms are more than half of all heavy atoms, speed-up due to smaller number of atoms is relatively insignificant, and replacement of local sampling by differentiation is the key underlying its efficiency. GSFE-Refinement is, to the best of our knowledge, the first end-to-end differentiable algorithm that takes fully trainable parameters and can generate a continuous dynamic trajectory similar to MD simulation. This unique property make it possible to realize physics based *ab initio* folding when properly trained for both unfolded and folded states, while likely to be significantly more efficient than MD simulation based folding.^31,32^ The competitive accuracy of our backbone and *C*_*β*_ representation on a par with mainstream all atom methods suggest that many body correlation captured by GSFE-Refinement are important. Simple backbone and *C*_*β*_ representation provides additional benefit of smoother FEL than all atom counterpart, its value in this regard is irreplaceable despite higher expected accuracy of all atom LFEL that is under development in our group. It is important to note that GSFE theory is one way of implementing LFEL for global free energy estimation and there might well be more elegant ways. While only gradient descent optimization is demonstrated in this work, AD is capable of calculating exact higher ordered derivatives and therefore exploration of higher ordered optimization algorithms in this scheme is certainly feasible and will be carried out in the future.

To cache or to compute intermediate results on the fly is a ubiquitous tradeoff in computation. As far as molecular free energy is concerned, we rely far more than necessary on computing by sampling, and much less on the inexpensive memory. “Local” in this work is conveniently defined as specific solvent units of each solute unit and its spatial coverage (i.e. cutoff) need to be determined to implement GSFE for construction of LFEL. The larger the “local” is, the more data and the larger neural networks are needed to cache corresponding LFEL, and the faster and more accurate computation will be achieved in subsequent free energy optimization. However, as strength of correlation decreases rapidly with distance, when “local” extends beyond certain level of correlation, the increase of data and training cost is likely to be unworthy and negative impact of noises may rise. Input for training LFEL may be of either computational or experimental origin. Apart from hardware consideration for the optimization, data availability and network architecture are of critical importance for training. In the case of PSR presented here, the size of the “local” is likely restricted either by the number of available high resolution experimental structures or by efficiency of network in extracting correlations. If significantly more surrounding residues were taken as the solvent of a target residue, reliable description of their spatial distributions would require more data and/or more efficient network. More investigations are necessary to understand relative importance of these two factors. When high quality experimental data is not available or not sufficient, a potentially feasible two-step strategy is to first utilize neural network FF to generate sufficiently large number of configurations for interested compositions at desired thermodynamic conditions (e.g. temperature and pressure). This step, if done properly, may realize sampling of reliable quantum mechanical accuracy. Secondly, these configurations may be subsequently utilized to construct LFEL for efficiently and reliably carrying out many free energy optimization tasks with near quantum accuracy and efficiency of traditional coarse grained methods. For proteins in particular, complete computation driven *ab initio* folding of proteins without learning from experimental structural information is potentially possible in this framework.

In equation 3, all local distributions seem be treated as independent. However, this is not the case in our implementation. Apart from direct long-range interactions, all mediated global correlations among overlapping local regions are embodied by the fact that they share the same coordinates. This constraint is exactly satisfied during all iterations as only one set of coordinates are utilized. At each cycle, each residue participating in LFEL of all its solvent units and its coordinates are updated as a result of compromise among their LFEL, result in a larger effective cutoff than defined by “local” in training of LFEL (see Fig. 1).

One great feature of our scheme is that coordinate update and transformation module is separated from LFEL. Therefore, future modification of neural network architecture for caching LFEL is flexible. For the specific task of PSR and its potential extension to *ab initio* protein folding, we indeed need such flexibility to advance from present backbone and *C*_*β*_ level LMLA treatment (Equation 3) to incorporate all side chain heavy atoms, local priors and direct long range interactions. Another advantage of GSFE is that direct control of each comprising unit is straight forward with well-defined physical interpretation as demonstrated by addition of local restraints (Equations 5 and 6). The scheme (Fig. 2) for optimizing molecular free energy on a computational graph is of general utility in soft matter modeling. It is also important that while brute force caching of LFEL is apparently specific for given constraint environmental conditions (e.g. temperature, pressure, composition), inclusion of these conditions within LFEL is possible and will be one interesting future research direction.

In light of the amazing success by AlphaFold2 for protein structure prediction, it is important to note that our algorithm is targeted to physical simulation and understanding of both static distributions (i.e. structure) and dynamic processes in complex molecular systems. This is in strong contrast to all mapping algorithms from sequences to structures through black-box modules as in AlphaFold2. LFEL is a new path for accelerating molecular simulations, its connection to enhanced sampling and coarse graining is detailed elsewhere.^18^

## Conclusions

In summary, we develop a novel scheme that maps molecular free energy optimization onto a computational graph through integrating GSFE theory, autodifferentiation and coordinate transformation. The key contribution is to replace expensive local sampling by differentiation w.r.t. LFEL, which is cached by fully trainable neural networks. Overlapping among many “local” region naturally maintains global mediated correlations by the simple fact that only one set of coordinates are utilized for all local regions in each iteration. As local sampling is repetitively carried out in essentially all present free energy simulations and consumes majority of computational resources, replacement of which by differentiation w.r.t. LFEL is expected to bring dramatic savings without loss of resolution. As an exemplary implementation of this scheme, we develop a backbone and *C*_*β*_ representation of PSR pipeline that relies solely on fully trainable description of LFEL for the first time. When compared with main-stream methods, this pipeline demonstrates competitive accuracy and is orders of magnitude more efficient. Further improvement in accuracy are expected with future incorporation of more input information and better representation of local prior term. Additionally, this is a general free energy optimization scheme for molecular systems of soft condensed matter. We hope our work stimulate more interest in formulation and methodology development in utilizing LFEL.

## Methods

### Introduction to local maximum likelihood approximation of GSFE

In GSFE, each comprising unit is both solute and solvent of its neighbors. For a n-residue protein with sequence *X* = {*x*_1_, *x*_2_, …, *x*_*n*_}, the free energy of a given structure is:

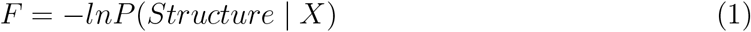

With Bayes formula:

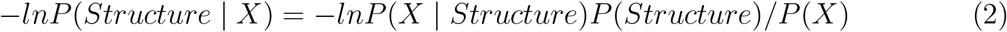

For a given sequence X, P(X) is a constant and is dropped. For maximum likelihood approximation, the prior term *P* (*Structure*) is ignored. Define *R*_*i*_(*X*_*i*_, *Y*_*i*_) as local structural regions within selected cutoff distance of *X*_*i*_ (*Y*_*i*_ being specific solvent of *X*_*i*_). With further local approximation we have:

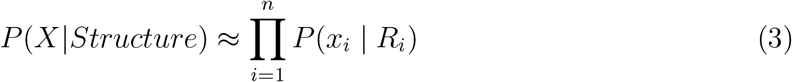

Equation 3 assumes all influence to each unit is included in its solvent *Y*_*i*_, this product of n local likelihood terms is the LMLA (local maximum likelihood approximation) of GSFE, which is the basis for our NN training in this work. Incorporation of local priors and direct long range interactions will be tackled in future.

### Training of local free energy landscape neural network

The same training/validation/test dataset as Ref^8^ is used for training LFEL. For each target residue, 22 neighboring residues are selected as its neighbors (6 upstream, 6 downstream sequentially adjacent residues in primary sequence and 10 non-adjacent ones). Features for each target residue include one-hot vectors representing identities of neighboring residues, residue pair distances (*C*_*α*_ −*C*_*α*_) between the target and its neighbor residues, and dihedral angles (*C*_*α*_, *C, N, C*_*β*_) indicating side chain orientations. Input for each comprising solvent unit of a LFEL includes a 22-dimensional one-hot vector, 6 sets of bond angle parameters (each angle *θ* is converted into sin*θ* and cos*θ*), and the *C*_*α*_ −*C*_*α*_ distance resulting in an input of (22 + 6 ∗ 2 + 1) ∗ 22 = 770 dimensions, the resulting LFEL is denoted as 770-LFEL. A four-layer feed-forward (770-512-512-512-21) network architecture is used (Fig. 2A) and trained for 30 epochs with a learning rate of 0.1.

### Refinement datasets

Four data sets are prepared to evaluate GSFE-Refinement. The first is 3DRobot data set. After removing structures having sequences of higher than 25% identity with training set, 36 of original 200 native structures^33^ remain, and 322 decoys for these 36 native structures are selected by random sampling according to refineD.^30^ The second is the 150-target RefineD test set.^30^ The third dataset includes 34 decoys available from CASP11 and 31 available from CASP12 respectively. The last data set is the 31 decoys out of 44 from CASP14 (We missed the earliest 13 targets due to late registration). CASP decoys are downloaded from CASP website (https://predictioncenter.org/download_area/).

### The refinement pipeline

As shown in Fig.2 B, a given starting structure is first striped of all side chain atoms other than *C*_*β*_. Cartesian coordinates of the remaining backbone (*C*_*α*_, *CO, N*) and *C*_*β*_ atoms are converted into internal coordinates, which is further converted back into cartesian coordinates for feature extraction. The NeRF (Natural Extension Reference Frame) algorithm (see support information for details) is used for coordinates transformation in both ways (see support information for details). The approximate free energy calculated by the forward pass through the neural network (which caches LFELs), plus some additional restraints (see below), constitute the total loss function. Derivatives of the total loss w.r.t. the input coordinates is calculated with back propagation of AD, and optimization is subsequently performed with simple gradient descent using the given learning rate and calculated derivatives. To maintain constraints of bond lengths and angles, only gradients of the loss function w.r.t. backbone dihedrals *ϕ* and *ψ* are saved, and derivatives w.r.t. other inputs are set to zero.

#### Loss function of the optimization process

The total loss is shown below:

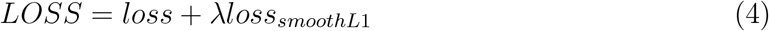

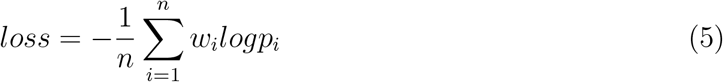

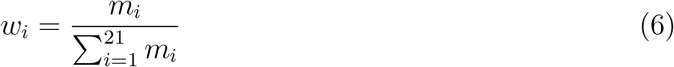

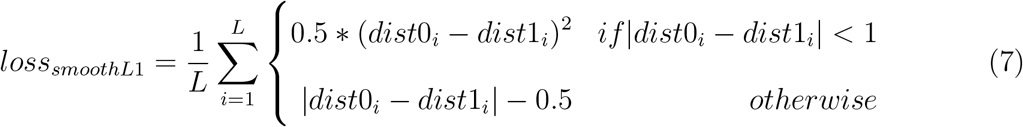

Here, *λ* is the coefficient of *loss*_*smoothL*1_, which is designed to limit the conformational search space during optimization. The larger the *λ*, the stronger the restraint. *w*_*i*_ is the AA (amino acid) site weight parameter, *n* is the protein chain length, *m* is the total number of AA types (m = 21), *L* is the number of neighboring AA around each target AA (L = 22), and *p*_*i*_ is the predicted likelihood for the specific solvent configuration given residue i, *dist*0_*i*_ and *dist*1_*i*_ are distances between the solute *C*_*a*_ and its *i*th solvent *C*_*a*_ atoms in the starting and updated structures. The total LOSS is iteratively minimized during the optimization. Learning rate specifies an effective step size for updating coordinates, its value is specified in *results*. It is important to note that learning rate in training process specifies magnitude for updating of neural network parameters, which caches LFEL.

## Supporting information

support information

## Acknowledgement

This work has been supported by the National Key Research and Development Program of China (2017YFB0702500) and by National Natural Science Foundation of China under grant number 31270758.

## Notes

### Competing Interest Statement

The authors have declared no competing interest.

### Summary of Updates

Significant modifications have been made to the manuscript. Title is changed to "Molecular free energy optimization on a computational graph". The previous emphasize the specific details of protein structure refinement, while this manuscript emphasize more on the general methodology for free energy optimization on a computational graph with protein structural refinement as a specific application. Latest results from CASP14 competition are included to strengthen the paper.

## References

(1) Free Energy Calculations; Springer: Berlin Heidelberg New York, 2007.

(2) Leman, J. K. et al. Macromolecular modeling and design in Rosetta: recent methods and frameworks. NATURE METHODS 2020, 17, 665–680.

(3) Roy, A., Kucukural, A., Zhang, Y. I-TASSER: a unified platform for automated protein structure and function prediction. NATURE PROTOCOLS 2010, 5, 725–738.

(4) Adiyaman, R., McGuffin, L. J. Methods for the Refinement of Protein Structure 3D Models. International Journal of Molecular Sciences 2019, 20.

(5) Khatib, F., Cooper, S., Tyka, M. D., Xu, K., Makedon, I., Popović, Z., Baker, D., Players, F. Algorithm discovery by protein folding game players. Proceedings of the National Academy of Sciences 2011, 108, 18949–18953.

(6) Zhang, J., Zhang, Y. A Novel Side-Chain Orientation Dependent Potential Derived from Random-Walk Reference State for Protein Fold Selection and Structure Prediction. PLOS ONE 2010, 5, 1–13.

(7) Uziela, K., Menéndez Hurtado, D., Shu, N., Wallner, B., Elofsson, A. ProQ3D: improved model quality assessments using deep learning. Bioinformatics 2017, 33, 1578–1580.

(8) Long, S., Tian, P. A simple neural network implementation of generalized solvation free energy for assessment of protein structural models. RSC Advances 2019, 9.

(9) Vanommeslaeghe, K., Hatcher, E., Acharya, C., Kundu, S., Zhong, S., Shim, J., Dar-ian, E., Guvench, O., Lopes, P., Vorobyov, I., Mackerell Jr., A. D. CHARMM general force field: A force field for drug-like molecules compatible with the CHARMM all-atom additive biological force fields. Journal of Computational Chemistry 2010, 31, 671–690.

(10) Alford, R. F. et al. The Rosetta All-Atom Energy Function for Macromolecular Modeling and Design. Journal of Chemical Theory and Computation 2017, 13, 3031–3048, PMID: 28430426.

(11) Nymeyer, H., García, A. E., Onuchic, J. N. Folding funnels and frustration in off-lattice minimalist protein landscapes. Proceedings of the National Academy of Sciences 1998, 95, 5921–5928.

(12) Ryckaert, J.-P., Ciccotti, G., Berendsen, H. J. Numerical integration of the cartesian equations of motion of a system with constraints: molecular dynamics of n-alkanes. Journal of Computational Physics 1977, 23, 327 – 341.

(13) Andersen, H.C. Rattle: A “velocity” version of the shake algorithm for molecular dynamics calculations. Journal of Computational Physics 1983, 52, 24–34.

(14) Miyamoto, S., Kollman, P. A. Settle: An analytical version of the SHAKE and RATTLE algorithm for rigid water models. Journal of Computational Chemistry 1992, 13, 952– 962.

(15) Dodd, L., Boone, T., Theodorou, D. A concerted rotation algorithm for atomistic Monte Carlo simulation of polymer melts and glasses. Molecular Physics 1993, 78, 961–996.

(16) Davis, I. W., Arendall, W. B., Richardson, D. C., Richardson, J. S. The Backrub Motion: How Protein Backbone Shrugs When a Sidechain Dances. Structure 2006, 14, 265 – 274.

(17) Gkeka, P. et al. Machine Learning Force Fields and Coarse-Grained Variables in Molecular Dynamics: Application to Materials and Biological Systems. Journal of Chemical Theory and Computation 2020, 16, 4757–4775, PMID: 32559068.

(18) Cao, X., Tian, P. “Dividing and Conquering” and “Caching” in Molecular Modeling. 2020.

(19) Werbos, P. J. Beyond regression: new tools for prediction and analysis in the behavioral sciences. Ph.D. thesis, Harvard University, 1974.

(20) Rumelhart, D., Ge, H., Rj, W. Learning representation by back-propagating errors. Nature 1986, 323, 533–536.

(21) Adam, Z. LGA: a method for finding 3D similarities in protein structures. Nuclc Acids Research 2003, 31, 3370–3374.

(22) Feig, M. Computational protein structure refinement: Almost there, yet still so far to go. Wiley Interdiplinary Reviews: Computational Molecular science 2017, 7, e1307.

(23) Qian, B., Raman, S., Das, R., Bradley, P., Mccoy, A. J., Read, R. J., Baker, D. High-resolution structure prediction and the crystallographic phase problem. Nature 2007, 450, 259–264.

(24) Nugent, T., Cozzetto, D., Jones, D. T. Evaluation of predictions in the CASP10 model refinement category. Proteins-structure Function Bioinformatics 2014, 82, 98–111.

(25) Parsons, J., Holmes, J. B., Rojas, J. M., Tsai, J., Strauss, C. E. M. Practical conversion from torsion space to Cartesian space for in silico protein synthesis. Journal of Computational Chemistry 2010, 26, 1063–1068.

(26) Ishitani, R., Terada, T., Shimizu, K. Refinement of comparative models of protein structure by using multicanonical molecular dynamics simulations. Molecular Simulation 2008, 34, 327–336.

(27) Cao, W., Terada, T., Nakamura, S., Shimizu, K. Refinement of Comparative-Modeling Structures by Multicanonical Molecular Dynamics. Genome Informatics International Conference on Genome Informatics 2011, 14, 484–485.

(28) Park, H., Seok, C. Refinement of unreliable local regions in template-based protein models. Proteins-structure Function Bioinformatics 2012, 80, 1974–1986.

(29) Zhang, J., Zhang, Y. A Novel Side-Chain Orientation Dependent Potential Derived from Random-Walk Reference State for Protein Fold Selection and Structure Prediction. Plos One 2010, 5, e15386.

(30) Debswapna, B. refineD: Improved protein structure refinement using machine learning based restrained relaxation. Bioinformatics 18.

(31) Shaw, D. E., Maragakis, P., Lindorff-Larsen, K., Piana, S., Dror, R. O., East-wood, M. P., Bank, J. A., Jumper, J. M., Salmon, J. K., Shan, Y., Wriggers, W. Atomic-Level Characterization of the Structural Dynamics of Proteins. SCIENCE 2010, 330, 341–346.

(32) Lane, T. J., Shukla, D., Beauchamp, K. A., Pande, V. S. To milliseconds and beyond: challenges in the simulation of protein folding. Current Opinion in Structural Biology 2013, 23, 58–65, Folding and binding / Protein-nucleic acid interactions.

(33) Haiyou, D., Ya, J., Yang, Z. 3DRobot: automated generation of diverse and well-packed protein structure decoys. Bioinformatics 2016, 32, 378–87.

