## Supplementary material for "Molecular free energy optimization on a computational graph": support information

Xiaoyong Cao[†] and Pu Tian[*,†,‡]

†*School of Life Sciences, Jilin University, Changchun, China 130012*

‡*School of Artificial Intelligence, Jilin University, Changchun, China 130012*

1.Coordinates transformation with NeRF (Natural Extension Reference Frame) algorithm:

$$\tilde{c_k} = [r_k cos(\theta_k), r_k cos(\psi_k) sin(\theta_k), r_k sin(\psi_k) cos(\theta_k)] \tag{1}$$

$$m_k = c_{k-1} - c_{k-2} \tag{2}$$

$$M_k = \widehat{m_k}, \widehat{m_k} \times \widehat{n_k}, \widehat{n_k} \tag{3}$$

$$c_k = M_k \tilde{c_k} + c_{k-1} \tag{4}$$

Here, $r_k$ is the bond length between atoms $k-1$ and $k$, $\theta_k$ is the bond angle composed of atoms $k-2$,$k-1$,$k$, and $\psi_k$ is the dihedral angle of $k-2$,$k-1$as the axis of rotation, $\widehat{m_k}$ the unit vector of $m_k$, $\times$ is the vector cross product, and $c_k$ is the cartesian coordinate of the updated $k$ atom.

Table S1: GSFE-Refinement best of top 5 results for 150-target refineD dataset

| ID | start GDT-HA | Δ GDT-HA | start RMSD | Δ RMSD |
|---|---|---|---|---|
| 101m_ | 0.6656 | 0.0146 | 3.9267 | -0.027 |
| 1a00A | 0.6418 | -0.0035 | 16.9914 | 0.0029 |
| 1a07A | 0.7405 | 0.0024 | 3.29 | -0.0044 |
| 1a5v_ | 0.5582 | 0.0034 | 1.8435 | -0.0214 |
| 1afqC | 0.5573 | 0.0078 | 6.2068 | -0.0098 |
| 1atlA | 0.64 | 0.0062 | 6.5628 | -0.0287 |
| 1awqA | 0.5732 | 0.003 | 9.9856 | -0.0217 |
| 1b0b_ | 0.6294 | -0.0017 | 1.837 | -0.0268 |
| 1b11A | 0.542 | -0.0022 | 12.6573 | -0.0301 |
| 1b1cA | 0.6627 | -0.0016 | 5.5089 | -0.0074 |
| 1b66A | 0.5707 | 0 | 0.8874 | -0.0338 |
| 1be9A | 0.5856 | 0.0024 | 5.3811 | -0.012 |
| 1bh5A | 0.531 | 0.0017 | 4.3604 | -0.0173 |
| 1bs4A | 0.5923 | 0.0015 | 5.9913 | -0.02 |
| 1bvqA | 0.5953 | 0.0072 | 4.5375 | -0.0135 |
| 1bzsA | 0.4152 | -0.0016 | 7.6959 | -0.022 |
| 1c9kB | 0.2056 | 0.0027 | 5.0313 | 0.0054 |
| 1ccwA | 0.6004 | 0.0036 | 2.454 | -0.0131 |
| 1cfvH | 0.6366 | 0 | 5.5613 | -0.0291 |
| 1cpq_ | 0.6647 | 0.0117 | 5.7355 | -0.0141 |
| 1cv8_ | 0.3613 | 0 | 3.5176 | -0.003 |
| 1d01D | 0.5685 | 0 | 6.0945 | -0.0368 |
| 1d0iA | 0.3883 | 0.0067 | 4.754 | -0.0419 |
| 1d1jC | 0.5328 | 0.0037 | 5.4926 | -0.0096 |
| 1d2uA | 0.3261 | 0.0027 | 4.994 | -0.0426 |

| | | | | |
|---|---|---|---|---|
| 1d7bA | 0.1041 | 0.0013 | 2.0833 | -0.0153 |
| 1d8uA | 0.4848 | 0.0016 | 2.8479 | -0.0197 |
| 1ddrA | 0.6368 | 0.0173 | 6.7574 | -0.0177 |
| 1df8A | 0.6229 | 0.0085 | 1.9512 | -0.0144 |
| 1ds7A | 0.5141 | 0.0077 | 2.1074 | -0.012 |
| 1dt1A | 0.3275 | 0.0078 | 1.794 | -0.0247 |
| 1dud_ | 0.5907 | 0.002 | 15.8571 | -0.047 |
| 1dveA | 0.4393 | 0.0093 | 1.1959 | -0.0107 |
| 1dw0A | 0.3281 | 0.0045 | 12.2524 | -0.0144 |
| 1dxpA | 0.2586 | 0.0043 | 11.966 | 0.008 |
| 1dzrA | 0.4522 | 0.0082 | 2.3676 | 0.0009 |
| 1e20A | 0.5176 | 0.0081 | 5.2297 | -0.0219 |
| 1e29A | 0.4222 | 0.0019 | 11.7805 | -0.0348 |
| 1e6iA | 0.7227 | 0 | 5.8339 | -0.0261 |
| 1ejbA | 0.6548 | 0.0014 | 5.8128 | -0.0672 |
| 1elkA | 0.5899 | 0.0065 | 3.8924 | -0.031 |
| 1elrA | 0.6074 | 0.0215 | 1.7917 | -0.0799 |
| 1elwA | 0.6859 | 0.0064 | 1.9984 | -0.0253 |
| 1eqmA | 0.5728 | 0.0031 | 11.3548 | -0.0094 |
| 1eumA | 0.7531 | 0.014 | 1.9421 | 0.0023 |
| 1ew4A | 0.5943 | 0.0024 | 1.857 | -0.015 |
| 1eyhA | 0.5434 | 0.0035 | 3.5525 | -0.0493 |
| 1f7lA | 0.5127 | 0.0106 | 1.9146 | -0.0194 |
| 1fdk_ | 0.2581 | -0.002 | 10.8596 | -0.0139 |
| 1ff3A | 0.5262 | 0.0048 | 2.1792 | -0.068 |
| 1fm0D | 0.7067 | 0 | 1.6562 | -0.0126 |
| 1g5tA | 0.4204 | 0.0032 | 6.5559 | -0.0181 |

| | | | | |
|---|---|---|---|---|
| 1gcvB | 0.6085 | 0.011 | 3.5266 | -0.081 |
| 1goyA | 0.3657 | 0 | 1.6183 | -0.0304 |
| 1gpqA | 0.4528 | 0.0019 | 5.4125 | -0.0131 |
| 1gxuA | 0.5881 | 0.0057 | 1.5192 | -0.0176 |
| 1gy7A | 0.6384 | 0.0124 | 4.0669 | -0.0019 |
| 1h4hA | 0.4782 | 0.0012 | 1.9931 | -0.0642 |
| 1h97A | 0.5408 | 0 | 1.4124 | 0.002 |
| 1hbg_ | 0.5986 | 0.0119 | 6.304 | -0.0138 |
| 1hdoA | 0.5671 | 0.0122 | 2.708 | -0.0462 |
| 1hlb_ | 0.5016 | 0.0143 | 1.4586 | -0.0285 |
| 1hnl_ | 0.2865 | 0.002 | 3.3574 | -0.0286 |
| 1i4sA | 0.449 | 0.0085 | 9.4846 | -0.0421 |
| 1j3wA | 0.5951 | 0.0131 | 2.7615 | 0.0016 |
| 1j77A | 0.4397 | 0.0063 | 3.8841 | -0.0585 |
| 1kafA | 0.2477 | 0.0023 | 3.6507 | -0.0028 |
| 1l9lA | 0.598 | -0.0034 | 2.6332 | 0.0024 |
| 1mk0A | 0.4459 | 0.0077 | 3.4147 | -0.0091 |
| 1mn8A | 0.2474 | 0 | 2.804 | -0.0494 |
| 1nlqA | 0.231 | 0.0047 | 11.5067 | -0.0315 |
| 1nwaA | 0.2411 | 0.0015 | 2.4345 | -0.0875 |
| 1o82A | 0.3036 | 0 | 3.9845 | -0.0081 |
| 1oh0A | 0.628 | 0.008 | 9.0235 | -0.0166 |
| 1ok0A | 0.1824 | 0 | 2.2832 | 0.008 |
| 1oohA | 0.4821 | 0.008 | 2.8792 | -0.0174 |
| 1ou8A | 0.4151 | 0.0024 | 5.0654 | -0.0272 |
| 1ow4A | 0.5479 | 0.0021 | 5.3922 | -0.0106 |
| 1q1fA | 0.625 | 0.0169 | 4.1534 | -0.0291 |

| | | | | |
|---|---|---|---|---|
| 1q4kA | 0.0788 | 0.0023 | 1.7395 | -0.0002 |
| 1rfyA | 0.618 | 0.0084 | 2.8979 | -0.0364 |
| 1sz7A | 0.4686 | 0.0047 | 8.7317 | -0.0081 |
| 1t1vA | 0.6156 | 0 | 8.2837 | -0.0289 |
| 1t3yA | 0.6221 | 0.0039 | 6.2022 | -0.0147 |
| 1t7dA | 0.0481 | 0 | 2.8585 | -0.0349 |
| 1tkuA | 0.0944 | 0 | 16.7339 | -0.1489 |
| 1tp6A | 0.4206 | 0.004 | 3.0161 | 0.0064 |
| 1tqgA | 0.7214 | 0.0048 | 1.1602 | -0.0205 |
| 1tugB | 0.4118 | 0 | 10.803 | -0.0408 |
| 1tzvA | 0.5177 | 0.0071 | 5.935 | -0.0097 |
| 1u2hA | 0.8021 | 0.0052 | 2.613 | 0.0013 |
| 1u55A | 0.4846 | 0.0126 | 2.4141 | -0.0309 |
| 1u84A | 0.3364 | 0.0031 | 2.9302 | -0.0031 |
| 1ufyA | 0.3417 | 0.0023 | 21.5158 | -0.0354 |
| 1ugiA | 0.1687 | 0 | 1.9191 | 0.0623 |
| 1usqA | 0.3291 | 0.0036 | 13.1353 | -0.004 |
| 1uuyA | 0.6491 | 0.0031 | 6.3911 | -0.0047 |
| 1uxzA | 0.6145 | -0.0019 | 4.1702 | -0.0192 |
| 1w53A | 0.503 | 0.0059 | 1.4101 | -0.0343 |
| 1wlzA | 0.3618 | 0.0029 | 19.0621 | -0.0217 |
| 1wmhA | 0.7289 | 0.003 | 9.198 | -0.0028 |
| 1wmhB | 0.5945 | 0 | 3.616 | -0.0027 |
| 1wpaA | 0.2477 | -0.0024 | 1.3425 | -0.0051 |
| 1wzdA | 0.4641 | 0.0024 | 1.5859 | 0.0076 |
| 1x6iA | 0.5337 | 0.0028 | 3.1087 | -0.0005 |
| 1x91A | 0.7232 | -0.0017 | 6.7188 | -0.0578 |

| | | | | |
|---|---|---|---|---|
| 1xa8A | 0.2296 | 0 | 13.2079 | -0.0558 |
| 1xmtA | 0.5579 | 0.0026 | 1.5011 | -0.0078 |
| 1xppA | 0.6237 | 0.0051 | 10.4144 | -0.0018 |
| 1y3dI | 0.3906 | 0.0078 | 5.3931 | -0.0326 |
| 1y9tA | 0.2023 | 0 | 1.5181 | -0.0034 |
| 1yd0A | 0.4494 | 0 | 2.3645 | -0.0371 |
| 1z3eB | 0.6607 | 0.0089 | 2.4512 | -0.0033 |
| 2b5aA | 0.776 | 0.013 | 6.9385 | -0.0475 |
| 2bwfA | 0.6461 | 0.0097 | 2.6113 | -0.005 |
| 2c92A | 0.6378 | 0.0034 | 3.1986 | 0.0076 |
| 2cb8A | 0.7674 | 0.0117 | 1.591 | -0.0114 |
| 2chhA | 0.1726 | 0.0022 | 2.2773 | 0.001 |
| 2ev1A | 0.1797 | 0 | 3.5855 | -0.0426 |
| 2fcwA | 0.4646 | 0.0024 | 3.4804 | -0.0177 |
| 2gkgA | 0.7049 | -0.0061 | 3.2566 | -0.0037 |
| 2grrB | 0.2102 | 0.0048 | 5.4023 | -0.0397 |
| 2h7zA | 0.53 | 0.0033 | 8.9504 | -0.0064 |
| 2hl0A | 0.5105 | 0 | 7.5891 | -0.0084 |
| 2ifrA | 0.1176 | 0.0013 | 5.0274 | -0.074 |
| 2ip6A | 0.7328 | 0.0057 | 5.6535 | -0.0169 |
| 2iu5A | 0.2891 | 0.0056 | 7.0724 | -0.023 |
| 2j8wA | 0.5547 | 0.0078 | 3.3315 | -0.0088 |
| 2jekA | 0.1411 | 0 | 1.632 | -0.0005 |
| 2nmlA | 0.205 | 0 | 2.7006 | -0.0192 |
| 2o37A | 0.5556 | 0.0061 | 8.8581 | -0.0286 |
| 2ofcA | 0.2234 | -0.0018 | 1.8955 | 0.0106 |
| 2oznA | 0.3985 | 0.0038 | 4.5841 | -0.0238 |

| | | | | |
|---|---|---|---|---|
| 2p6wA | 0.2124 | 0.006 | 8.5653 | -0.0531 |
| 2pmrA | 0.727 | 0.0033 | 2.5528 | -0.0423 |
| 2pv2A | 0.6796 | -0.0024 | 4.5033 | -0.0038 |
| 2pvbA | 0.4042 | 0 | 2.0095 | 0.0095 |
| 2rb8A | 0.7957 | 0.0027 | 1.9139 | -0.0008 |
| 2rk3A | 0.5963 | -0.0027 | 2.3792 | -0.0369 |
| 2v2pA | 0.8 | 0.0132 | 1.1419 | 0.0117 |
| 2v33A | 0.1978 | 0.0027 | 2.3929 | -0.011 |
| 2vyyA | 0.4727 | 0.0046 | 7.0412 | -0.01 |
| 2zs0D | 0.6793 | 0.0035 | 8.395 | -0.0025 |
| 3bfoA | 0.7118 | 0.0029 | 2.3744 | 0.0006 |
| 3bqpA | 0.5969 | 0 | 16.4106 | -0.0442 |
| 3bqsA | 0.2824 | 0 | 1.9198 | 0.0053 |
| 3by4A | 0.3939 | 0.0014 | 1.9598 | -0.0237 |
| 3c7mA | 0.4526 | 0.0077 | 11.8375 | -0.0006 |
| 3cjsB | 0.5729 | 0.0174 | 2.2228 | -0.0061 |
| 3d9nA | 0.5543 | 0.0091 | 1.5299 | -0.0188 |
| mean | - | 0.0041 | - | -0.0203 |

Table S2: GSFE-Refinement top 1 results for 150-target refineD dataset

| ID | start GDT-HA | Δ GDT-HA | start RMSD | Δ RMSD |
|---|---|---|---|---|
| 101m_ | 0.6656 | 0.0113 | 3.9267 | -0.0189 |
| 1a00A | 0.6418 | 0 | 16.9914 | -0.0037 |
| 1a07A | 0.7405 | 0 | 3.29 | 0.0074 |
| 1a5v_ | 0.5582 | 0 | 1.8435 | -0.013 |
| 1afqC | 0.5573 | -0.0026 | 6.2068 | 0.0095 |
| 1atlA | 0.64 | 0.0012 | 6.5628 | -0.0081 |

| | | | | |
|---|---|---|---|---|
| 1awqA | 0.5732 | -0.0031 | 9.9856 | -0.0013 |
| 1b0b_ | 0.6294 | -0.0159 | 1.837 | 0.0052 |
| 1b11A | 0.542 | 0 | 12.6573 | -0.0088 |
| 1b1cA | 0.6627 | 0 | 5.5089 | 0.0151 |
| 1b66A | 0.5707 | 0.0036 | 0.8874 | 0.0153 |
| 1be9A | 0.5856 | 0.0024 | 5.3811 | -0.0146 |
| 1bh5A | 0.531 | -0.0016 | 4.3604 | -0.0021 |
| 1bs4A | 0.5923 | 0 | 5.9913 | 0.01 |
| 1bvqA | 0.5953 | 0.0018 | 4.5375 | -0.0093 |
| 1bzsA | 0.4152 | 0 | 7.6959 | -0.0079 |
| 1c9kB | 0.2056 | -0.0014 | 5.0313 | 0.0296 |
| 1ccwA | 0.6004 | -0.0055 | 2.454 | 0.0077 |
| 1cfvH | 0.6366 | -0.0021 | 5.5613 | -0.0122 |
| 1cpq_ | 0.6647 | 0.0058 | 5.7355 | -0.0191 |
| 1cv8_ | 0.3613 | -0.0029 | 3.5176 | 0.0071 |
| 1d01D | 0.5685 | 0 | 6.0945 | 0.017 |
| 1d0iA | 0.3883 | 0 | 4.754 | 0.001 |
| 1d1jC | 0.5328 | 0.0055 | 5.4926 | -0.0115 |
| 1d2uA | 0.3261 | -0.0014 | 4.994 | -0.0026 |
| 1d7bA | 0.1041 | 0.0013 | 2.0833 | 0.0004 |
| 1d8uA | 0.4848 | 0.0031 | 2.8479 | -0.0081 |
| 1ddrA | 0.6368 | 0.0047 | 6.7574 | -0.0026 |
| 1df8A | 0.6229 | 0.0085 | 1.9512 | -0.0011 |
| 1ds7A | 0.5141 | 0 | 2.1074 | 0.0058 |
| 1dt1A | 0.3275 | 0.0058 | 1.794 | 0.0076 |
| 1dud_ | 0.5907 | 0.0061 | 15.8571 | -0.005 |
| 1dveA | 0.4393 | -0.0024 | 1.1959 | 0.0109 |

| | | | | |
|---|---|---|---|---|
| 1dw0A | 0.3281 | -0.0044 | 12.2524 | 0.0008 |
| 1dxpA | 0.2586 | 0.0014 | 11.966 | 0.0242 |
| 1dzrA | 0.4522 | -0.0014 | 2.3676 | -0.0008 |
| 1e20A | 0.5176 | 0 | 5.2297 | -0.0076 |
| 1e29A | 0.4222 | 0 | 11.7805 | -0.0003 |
| 1e6iA | 0.7227 | -0.0113 | 5.8339 | 0.0343 |
| 1ejbA | 0.6548 | -0.006 | 5.8128 | -0.0151 |
| 1elkA | 0.5899 | 0.0016 | 3.8924 | -0.0276 |
| 1elrA | 0.6074 | 0.0156 | 1.7917 | -0.0894 |
| 1elwA | 0.6859 | -0.0085 | 1.9984 | 0.0374 |
| 1eqmA | 0.5728 | 0.0016 | 11.3548 | 0.0197 |
| 1eumA | 0.7531 | 0 | 1.9421 | -0.0049 |
| 1ew4A | 0.5943 | 0.0048 | 1.857 | -0.0065 |
| 1eyhA | 0.5434 | -0.0035 | 3.5525 | -0.0851 |
| 1f7lA | 0.5127 | 0 | 1.9146 | -0.0004 |
| 1fdk_ | 0.2581 | -0.004 | 10.8596 | 0.0076 |
| 1ff3A | 0.5262 | -0.0024 | 2.1792 | 0.0105 |
| 1fm0D | 0.7067 | -0.0034 | 1.6562 | -0.0157 |
| 1g5tA | 0.4204 | 0 | 6.5559 | 0.0004 |
| 1gcvB | 0.6085 | 0.0018 | 3.5266 | -0.009 |
| 1goyA | 0.3657 | 0.0024 | 1.6183 | 0.0001 |
| 1gpqA | 0.4528 | 0.0039 | 5.4125 | -0.0012 |
| 1gxuA | 0.5881 | 0.0057 | 1.5192 | -0.0129 |
| 1gy7A | 0.6384 | 0.0062 | 4.0669 | 0.012 |
| 1h4hA | 0.4782 | -0.0025 | 1.9931 | 0.0015 |
| 1h97A | 0.5408 | 0 | 1.4124 | -0.0064 |
| 1hbg_ | 0.5986 | 0.0102 | 6.304 | -0.0236 |

| | | | | |
|---|---|---|---|---|
| 1hdoA | 0.5671 | 0.0024 | 2.708 | -0.0048 |
| 1hlb_ | 0.5016 | 0 | 1.4586 | -0.0121 |
| 1hnl_ | 0.2865 | -0.0019 | 3.3574 | 0.0107 |
| 1i4sA | 0.449 | 0.0068 | 9.4846 | -0.0085 |
| 1j3wA | 0.5951 | 0.0094 | 2.7615 | 0.0037 |
| 1j77A | 0.4397 | 0.005 | 3.8841 | 0.0023 |
| 1kafA | 0.2477 | 0.0046 | 3.6507 | 0.0114 |
| 1l9lA | 0.598 | -0.0034 | 2.6332 | 0.0025 |
| 1mk0A | 0.4459 | -0.0052 | 3.4147 | 0.0002 |
| 1mn8A | 0.2474 | -0.0027 | 2.804 | -0.0414 |
| 1nlqA | 0.231 | 0.0023 | 11.5067 | -0.0044 |
| 1nwaA | 0.2411 | 0.0015 | 2.4345 | -0.0059 |
| 1o82A | 0.3036 | 0 | 3.9845 | -0.0014 |
| 1oh0A | 0.628 | 0.008 | 9.0235 | 0.0107 |
| 1ok0A | 0.1824 | -0.0067 | 2.2832 | 0.0202 |
| 1oohA | 0.4821 | 0.004 | 2.8792 | -0.0013 |
| 1ou8A | 0.4151 | 0 | 5.0654 | -0.0295 |
| 1ow4A | 0.5479 | 0.0021 | 5.3922 | -0.0196 |
| 1q1fA | 0.625 | 0.0118 | 4.1534 | -0.0203 |
| 1q4kA | 0.0788 | 0.0012 | 1.7395 | 0.018 |
| 1rfyA | 0.618 | -0.0028 | 2.8979 | -0.001 |
| 1sz7A | 0.4686 | -0.0016 | 8.7317 | 0.014 |
| 1t1vA | 0.6156 | 0.0027 | 8.2837 | -0.0257 |
| 1t3yA | 0.6221 | 0.0039 | 6.2022 | -0.0084 |
| 1t7dA | 0.0481 | 0 | 2.8585 | -0.0319 |
| 1tkuA | 0.0944 | 0 | 16.7339 | -0.0069 |
| 1tp6A | 0.4206 | 0.002 | 3.0161 | 0.0289 |

| | | | | |
|---|---|---|---|---|
| 1tqgA | 0.7214 | 0.0048 | 1.1602 | -0.0117 |
| 1tugB | 0.4118 | 0 | 10.803 | 0.0133 |
| 1tzvA | 0.5177 | 0 | 5.935 | -0.0118 |
| 1u2hA | 0.8021 | 0 | 2.613 | 0.0007 |
| 1u55A | 0.4846 | 0.0042 | 2.4141 | 0.0006 |
| 1u84A | 0.3364 | 0 | 2.9302 | -0.0032 |
| 1ufyA | 0.3417 | 0.0023 | 21.5158 | 0.0061 |
| 1ugiA | 0.1687 | -0.003 | 1.9191 | 0.0474 |
| 1usqA | 0.3291 | 0.0036 | 13.1353 | 0.0028 |
| 1uuyA | 0.6491 | -0.0031 | 6.3911 | 0.0039 |
| 1uxzA | 0.6145 | 0 | 4.1702 | -0.0058 |
| 1w53A | 0.503 | 0.003 | 1.4101 | -0.0128 |
| 1wlzA | 0.3618 | -0.003 | 19.0621 | -0.0336 |
| 1wmhA | 0.7289 | 0 | 9.198 | 0.0112 |
| 1wmhB | 0.5945 | -0.0061 | 3.616 | 0.0024 |
| 1wpaA | 0.2477 | -0.0047 | 1.3425 | 0.1414 |
| 1wzdA | 0.4641 | 0 | 1.5859 | 0.0075 |
| 1x6iA | 0.5337 | 0 | 3.1087 | 0.0018 |
| 1x91A | 0.7232 | 0.0016 | 6.7188 | -0.0358 |
| 1xa8A | 0.2296 | 0.0025 | 13.2079 | -0.0074 |
| 1xmtA | 0.5579 | 0.0053 | 1.5011 | -0.0077 |
| 1xppA | 0.6237 | -0.0126 | 10.4144 | 0.1296 |
| 1y3dI | 0.3906 | 0.0039 | 5.3931 | -0.0249 |
| 1y9tA | 0.2023 | 0.0022 | 1.5181 | 0.0296 |
| 1yd0A | 0.4494 | -0.0056 | 2.3645 | -0.0069 |
| 1z3eB | 0.6607 | 0.0089 | 2.4512 | -0.0244 |
| 2b5aA | 0.776 | 0.0065 | 6.9385 | -0.004 |

| | | | | |
|---|---|---|---|---|
| 2bwfA | 0.6461 | 0.0033 | 2.6113 | 0.014 |
| 2c92A | 0.6378 | -0.0017 | 3.1986 | 0.0072 |
| 2cb8A | 0.7674 | 0 | 1.591 | -0.0038 |
| 2chhA | 0.1726 | -0.0022 | 2.2773 | 0.0297 |
| 2ev1A | 0.1797 | 0.0014 | 3.5855 | -0.008 |
| 2fcwA | 0.4646 | 0 | 3.4804 | -0.0136 |
| 2gkgA | 0.7049 | -0.0061 | 3.2566 | -0.0029 |
| 2grrB | 0.2102 | 0.0048 | 5.4023 | -0.0127 |
| 2h7zA | 0.53 | -0.0067 | 8.9504 | 0.0084 |
| 2hl0A | 0.5105 | 0 | 7.5891 | 0.012 |
| 2ifrA | 0.1176 | 0.0013 | 5.0274 | 0.0034 |
| 2ip6A | 0.7328 | 0.0028 | 5.6535 | 0.0104 |
| 2iu5A | 0.2891 | 0.0014 | 7.0724 | -0.0131 |
| 2j8wA | 0.5547 | 0.0019 | 3.3315 | -0.0211 |
| 2jekA | 0.1411 | -0.0018 | 1.632 | -0.0091 |
| 2nmlA | 0.205 | 0.0025 | 2.7006 | 0.0044 |
| 2o37A | 0.5556 | 0.0123 | 8.8581 | -0.0356 |
| 2ofcA | 0.2234 | -0.0018 | 1.8955 | 0.0123 |
| 2oznA | 0.3985 | -0.0019 | 4.5841 | -0.002 |
| 2p6wA | 0.2124 | -0.0012 | 8.5653 | 0.0246 |
| 2pmrA | 0.727 | 0 | 2.5528 | 0.0046 |
| 2pv2A | 0.6796 | 0.0073 | 4.5033 | -0.0095 |
| 2pvbA | 0.4042 | -0.0023 | 2.0095 | 0.0169 |
| 2rb8A | 0.7957 | 0 | 1.9139 | -0.0047 |
| 2rk3A | 0.5963 | -0.0027 | 2.3792 | 0.0209 |
| 2v2pA | 0.8 | 0.0029 | 1.1419 | 0.0013 |
| 2v33A | 0.1978 | 0.0082 | 2.3929 | -0.0045 |

| | | | | |
|---|---|---|---|---|
| 2vyyA | 0.4727 | 0.0023 | 7.0412 | 0.0048 |
| 2zs0D | 0.6793 | -0.0034 | 8.395 | -0.0017 |
| 3bfoA | 0.7118 | 0 | 2.3744 | 0.0006 |
| 3bqpA | 0.5969 | 0.0031 | 16.4106 | 0.0045 |
| 3bqsA | 0.2824 | 0 | 1.9198 | -0.0101 |
| 3by4A | 0.3939 | -0.0015 | 1.9598 | 0.0131 |
| 3c7mA | 0.4526 | 0 | 11.8375 | -0.005 |
| 3cjsB | 0.5729 | 0.007 | 2.2228 | -0.0123 |
| 3d9nA | 0.5543 | 0.0055 | 1.5299 | -0.0182 |
| mean | - | 0.0008 | - | -0.0003 |

Table S3: GSFE-Refinement best of top 5 results for CASP12 dataset

| ID | start GDT-HA | Δ GDT-HA | start RMSD | Δ RMSD |
|---|---|---|---|---|
| TR520 | 0.581 | -0.0031 | 3.9267 | -0.0012 |
| TR594 | 0.3427 | 0 | 16.9914 | -0.0057 |
| TR862 | 0.4032 | 0.0027 | 3.29 | -0.0449 |
| TR866 | 0.55 | 0 | 1.8435 | -0.0181 |
| TR868 | 0.5733 | 0.0043 | 6.2068 | -0.006 |
| TR869 | 0.2885 | 0 | 6.5628 | -0.004 |
| TR870 | 0.2276 | 0.0021 | 9.9856 | -0.0051 |
| TR872 | 0.5682 | 0 | 1.837 | -0.021 |
| TR877 | 0.4894 | 0.0053 | 12.6573 | -0.0319 |
| TR879 | 0.6352 | -0.0079 | 5.5089 | -0.0899 |
| TR882 | 0.6899 | 0.0031 | 0.8874 | -0.034 |
| TR884 | 0.4401 | 0.0036 | 5.3811 | -0.0155 |
| TR885 | 0.7412 | 0 | 4.3604 | -0.0135 |
| TR891 | 0.7567 | 0.0022 | 5.9913 | -0.0003 |
| TR893 | 0.6908 | 0.0015 | 4.5375 | -0.0094 |
| TR894 | 0.5463 | 0 | 7.6959 | -0.0095 |
| TR895 | 0.5146 | 0.0042 | 5.0313 | -0.0721 |
| TR896 | 0.468 | 0.0029 | 2.454 | -0.0065 |
| TR898 | 0.2524 | 0.0023 | 5.5613 | -0.0148 |
| TR905 | 0.3244 | 0.0051 | 5.7355 | -0.0421 |
| TR909 | 0.4264 | -0.003 | 3.5176 | -0.0336 |
| TR913 | 0.4534 | -0.003 | 6.0945 | -0.0429 |
| TR917 | 0.6535 | 0.0211 | 4.754 | -0.0668 |
| TR920 | 0.6039 | 0.0023 | 5.4926 | -0.0238 |
| TR921 | 0.4801 | 0.0036 | 4.994 | -0.0093 |
| TR922 | 0.6791 | 0.0067 | 2.0833 | -0.0078 |
| TR928 | 0.4274 | 0.0059 | 2.8479 | -0.0215 |
| TR944 | 0.5603 | 0.002 | 6.7574 | -0.0275 |
| TR945 | 0.4113 | -0.002 | 1.9512 | -0.0604 |
| TR947 | 0.5157 | 0.0143 | 2.1074 | -0.0456 |
| TR948 | 0.5956 | 0.0084 | 1.794 | -0.0105 |
| mean | - | 0.0027 | - | -0.0146 |

Table S4: GSFE-Refinement top 1 results for CASP12 dataset

| ID | start GDT-HA | $\Delta$ GDT-HA | start RMSD | $\Delta$ RMSD |
|---|---|---|---|---|
| TR520 | 0.581 | -0.0062 | 3.9267 | -0.0012 |
| TR594 | 0.3427 | 0 | 16.9914 | 0.0046 |
| TR862 | 0.4032 | 0 | 3.29 | 0.0209 |
| TR866 | 0.55 | -0.0065 | 1.8435 | 0.0012 |
| TR868 | 0.5733 | 0.0043 | 6.2068 | 0.0029 |
| TR869 | 0.2885 | 0 | 6.5628 | -0.0035 |
| TR870 | 0.2276 | -0.002 | 9.9856 | 0.0198 |
| TR872 | 0.5682 | 0 | 1.837 | 0.0052 |
| TR877 | 0.4894 | -0.0052 | 12.6573 | 0.007 |
| TR879 | 0.6352 | -0.0079 | 5.5089 | 0.0764 |
| TR882 | 0.6899 | 0 | 0.8874 | 0.0154 |
| TR884 | 0.4401 | 0.0036 | 5.3811 | -0.0143 |
| TR885 | 0.7412 | -0.0044 | 4.3604 | -0.0135 |
| TR891 | 0.7567 | 0.0022 | 5.9913 | -0.0003 |
| TR893 | 0.6908 | 0 | 4.5375 | 0.0084 |
| TR894 | 0.5463 | -0.0046 | 7.6959 | 0.0031 |
| TR895 | 0.5146 | -0.0063 | 5.0313 | 0.0438 |
| TR896 | 0.468 | 0.0029 | 2.454 | 0.0012 |
| TR898 | 0.2524 | 0 | 5.5613 | -0.0001 |
| TR905 | 0.3244 | 0.0051 | 5.7355 | -0.0421 |
| TR909 | 0.4264 | -0.003 | 3.5176 | 0.015 |
| TR913 | 0.4534 | -0.0089 | 6.0945 | -0.0429 |
| TR917 | 0.6535 | -0.0269 | 4.754 | 0.0792 |
| TR920 | 0.6039 | 0.0023 | 5.4926 | -0.0213 |
| TR921 | 0.4801 | -0.0091 | 4.994 | 0.0197 |
| TR922 | 0.6791 | 0.0033 | 2.0833 | 0.0039 |
| TR928 | 0.4274 | -0.0146 | 2.8479 | 0.0557 |
| TR944 | 0.5603 | -0.001 | 6.7574 | -0.0243 |
| TR945 | 0.4113 | -0.0206 | 1.9512 | 0.0684 |
| TR947 | 0.5157 | -0.0086 | 2.1074 | -0.0456 |
| TR948 | 0.5956 | -0.0033 | 1.794 | 0.0136 |
| mean | - | -0.0037 | - | 0.0083 |

Table S5: GSFE-Refinement best of top 5 results for CASP11 dataset

| ID | start GDT-HA | Δ GDT-HA | start RMSD | Δ RMSD |
|---|---|---|---|---|
| TR217 | 0.644 | 0.0072 | 3.9267 | -0.0128 |
| TR228 | 0.5476 | 0 | 16.9914 | -0.0325 |
| TR274 | 0.2896 | 0.0041 | 3.29 | -0.0186 |
| TR280 | 0.5938 | 0 | 1.8435 | -0.0266 |
| TR283 | 0.4119 | 0.0128 | 6.2068 | -0.0038 |
| TR759 | 0.4395 | 0 | 6.5628 | -0.0176 |
| TR762 | 0.7043 | 0.001 | 9.9856 | -0.0168 |
| TR765 | 0.5789 | 0.0033 | 1.837 | -0.0136 |
| TR768 | 0.6381 | 0.0035 | 12.6573 | -0.0175 |
| TR769 | 0.5851 | 0 | 5.5089 | 0.0079 |
| TR772 | 0.524 | 0 | 0.8874 | -0.0062 |
| TR774 | 0.3767 | 0.005 | 5.3811 | -0.0105 |
| TR776 | 0.6279 | 0.0148 | 4.3604 | -0.0103 |
| TR780 | 0.5395 | 0 | 5.9913 | -0.0083 |
| TR782 | 0.6477 | 0.0046 | 4.5375 | -0.0207 |
| TR783 | 0.573 | 0 | 7.6959 | -0.0184 |
| TR786 | 0.4781 | 0 | 5.0313 | -0.0155 |
| TR792 | 0.5813 | 0.0062 | 2.454 | -0.0145 |
| TR795 | 0.5864 | 0 | 5.5613 | 0.0078 |
| TR803 | 0.3321 | 0.0037 | 5.7355 | 0.005 |
| TR810 | 0.54 | -0.0044 | 3.5176 | -0.0269 |
| TR811 | 0.7331 | -0.002 | 6.0945 | 0.0023 |
| TR816 | 0.5221 | -0.0037 | 4.754 | -0.0143 |
| TR821 | 0.4833 | 0.0128 | 5.4926 | -0.0059 |
| TR822 | 0.3026 | 0.0044 | 4.994 | -0.0047 |
| TR827 | 0.3394 | 0.009 | 2.0833 | -0.048 |
| TR828 | 0.4881 | 0.0059 | 2.8479 | -0.0255 |
| TR829 | 0.5037 | 0 | 6.7574 | -0.0065 |
| TR833 | 0.6134 | -0.0023 | 1.9512 | -0.0206 |
| TR837 | 0.4339 | 0 | 2.1074 | -0.0485 |
| TR848 | 0.5815 | -0.0018 | 1.794 | -0.0267 |
| TR854 | 0.5857 | 0.0036 | 15.8571 | -0.0189 |
| TR856 | 0.6148 | 0 | 1.1959 | -0.0012 |
| TR857 | 0.3307 | 0.0052 | 12.2524 | -0.0066 |
| mean | - | 0.0027 | - | -0.0145 |

Table S6: GSFE-Refinement top 1 results for CASP11 dataset

| ID | start GDT-HA | Δ GDT-HA | start RMSD | Δ RMSD |
|---|---|---|---|---|
| TR217 | 0.644 | 0.006 | 3.9267 | -0.0096 |
| TR228 | 0.5476 | -0.0059 | 16.9914 | -0.0356 |
| TR274 | 0.2896 | 0 | 3.29 | 0.0016 |
| TR280 | 0.5938 | 0.0026 | 1.8435 | -0.0213 |
| TR283 | 0.4119 | 0.0032 | 6.2068 | -0.0031 |
| TR759 | 0.4395 | 0.004 | 6.5628 | 0.0009 |
| TR762 | 0.7043 | 0.0039 | 9.9856 | -0.0057 |
| TR765 | 0.5789 | 0.0033 | 1.837 | -0.0067 |
| TR768 | 0.6381 | 0.0018 | 12.6573 | -0.0059 |
| TR769 | 0.5851 | 0.0025 | 5.5089 | -0.0004 |
| TR772 | 0.524 | -0.0013 | 0.8874 | 0.002 |
| TR774 | 0.3767 | 0.0033 | 5.3811 | -0.0065 |
| TR776 | 0.6279 | 0.0045 | 4.3604 | -0.0026 |
| TR780 | 0.5395 | 0.0052 | 5.9913 | 0.0069 |
| TR782 | 0.6477 | -0.0022 | 4.5375 | -0.0067 |
| TR783 | 0.573 | 0 | 7.6959 | 0.0106 |
| TR786 | 0.4781 | -0.0034 | 5.0313 | 0.0006 |
| TR792 | 0.5813 | 0.0031 | 2.454 | -0.0143 |
| TR795 | 0.5864 | -0.0055 | 5.5613 | 0.009 |
| TR803 | 0.3321 | 0.0019 | 5.7355 | 0.02 |
| TR810 | 0.54 | 0.0011 | 3.5176 | -0.0113 |
| TR811 | 0.7331 | 0.001 | 6.0945 | 0.0005 |
| TR816 | 0.5221 | -0.0074 | 4.754 | -0.0018 |
| TR821 | 0.4833 | -0.0019 | 5.4926 | 0.0031 |
| TR822 | 0.3026 | 0.0022 | 4.994 | -0.0089 |
| TR827 | 0.3394 | 0.0065 | 2.0833 | -0.0456 |
| TR828 | 0.4881 | 0.0059 | 2.8479 | -0.009 |
| TR829 | 0.5037 | 0 | 6.7574 | 0.0016 |
| TR833 | 0.6134 | -0.0046 | 1.9512 | 0.0015 |
| TR837 | 0.4339 | 0.0041 | 2.1074 | -0.0289 |
| TR848 | 0.5815 | 0 | 1.794 | -0.0016 |
| TR854 | 0.5857 | 0.0107 | 15.8571 | -0.0151 |
| TR856 | 0.6148 | 0 | 1.1959 | -0.008 |
| TR857 | 0.3307 | 0.0026 | 12.2524 | -0.0018 |
| mean | - | 0.0014 | - | -0.0056 |

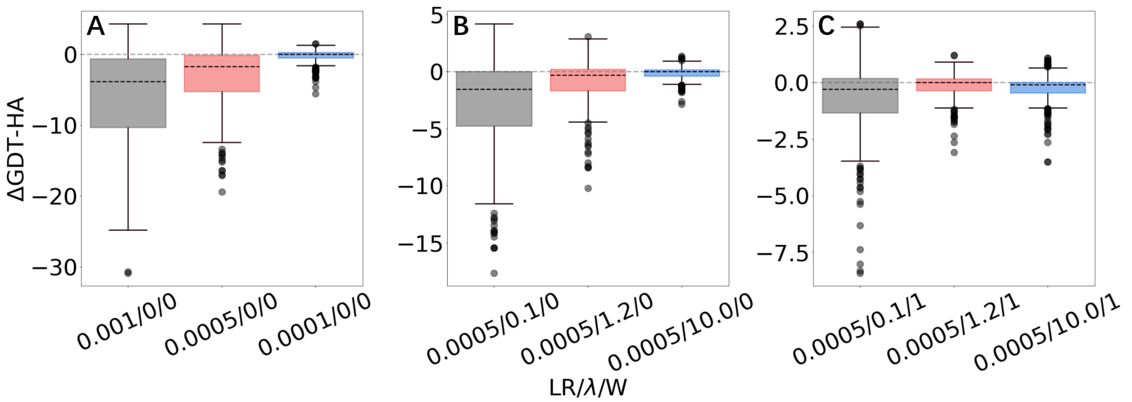


Figure S1: ΔGDT-HA of top 1 models by GFSE-Refinement for various /LR/λ/W combinations on 3DRobot dataset

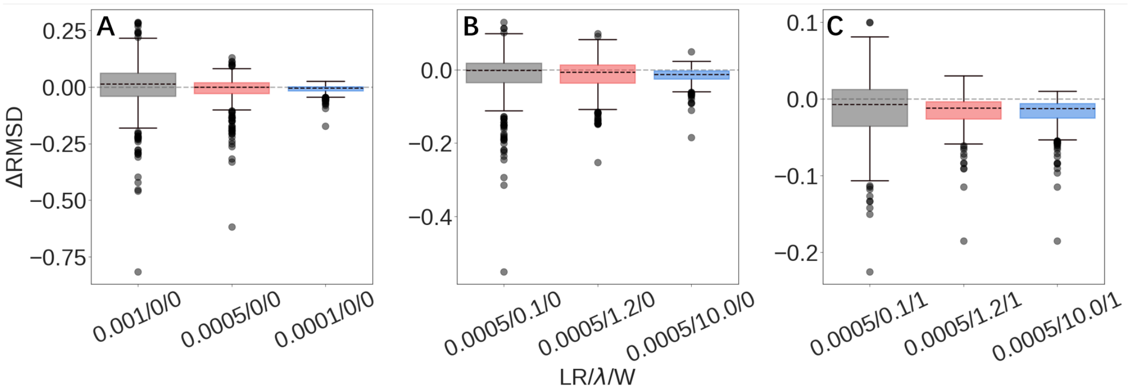


Figure S2: ΔRMSD of best of top 5 models by GFSE-Refinement for various /LR/λ/W combinations on 3DRobot dataset

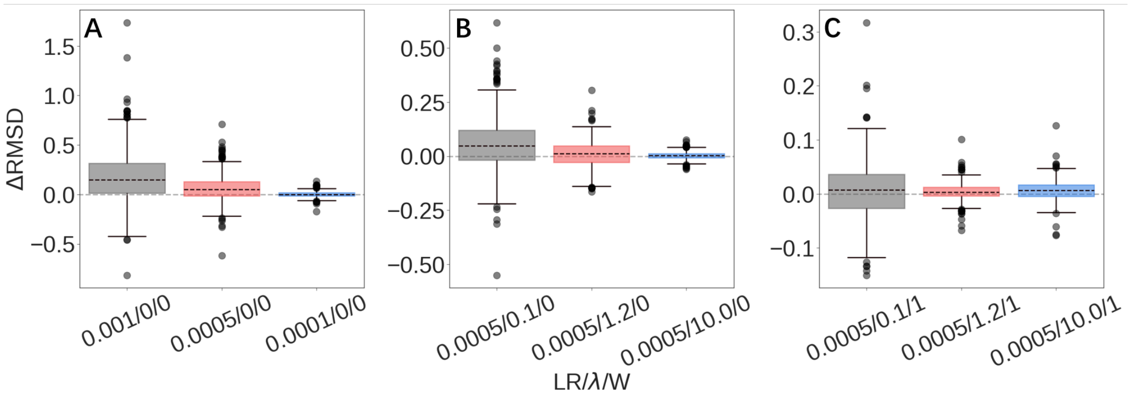


Figure S3: ΔRMSD of top 1 models by GFSE-Refinement for various /LR/$\lambda$/W combinations on 3DRobot dataset

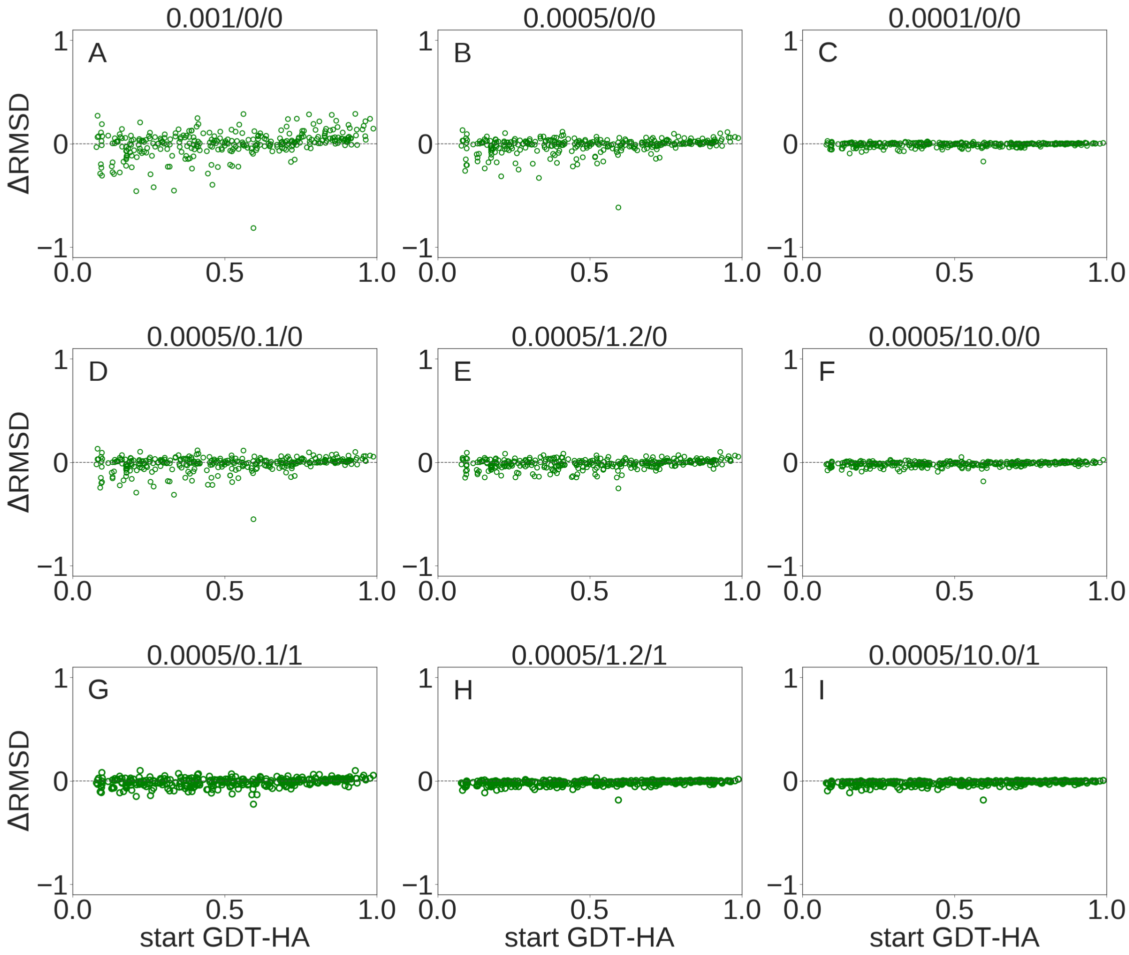


Figure S4: Scatter plots of ΔRMSD as a function of start GDT-HA for best of top 5 models by GSFE-Refinement for various /LR/$\lambda$/W combinations on 3DRobot dataset

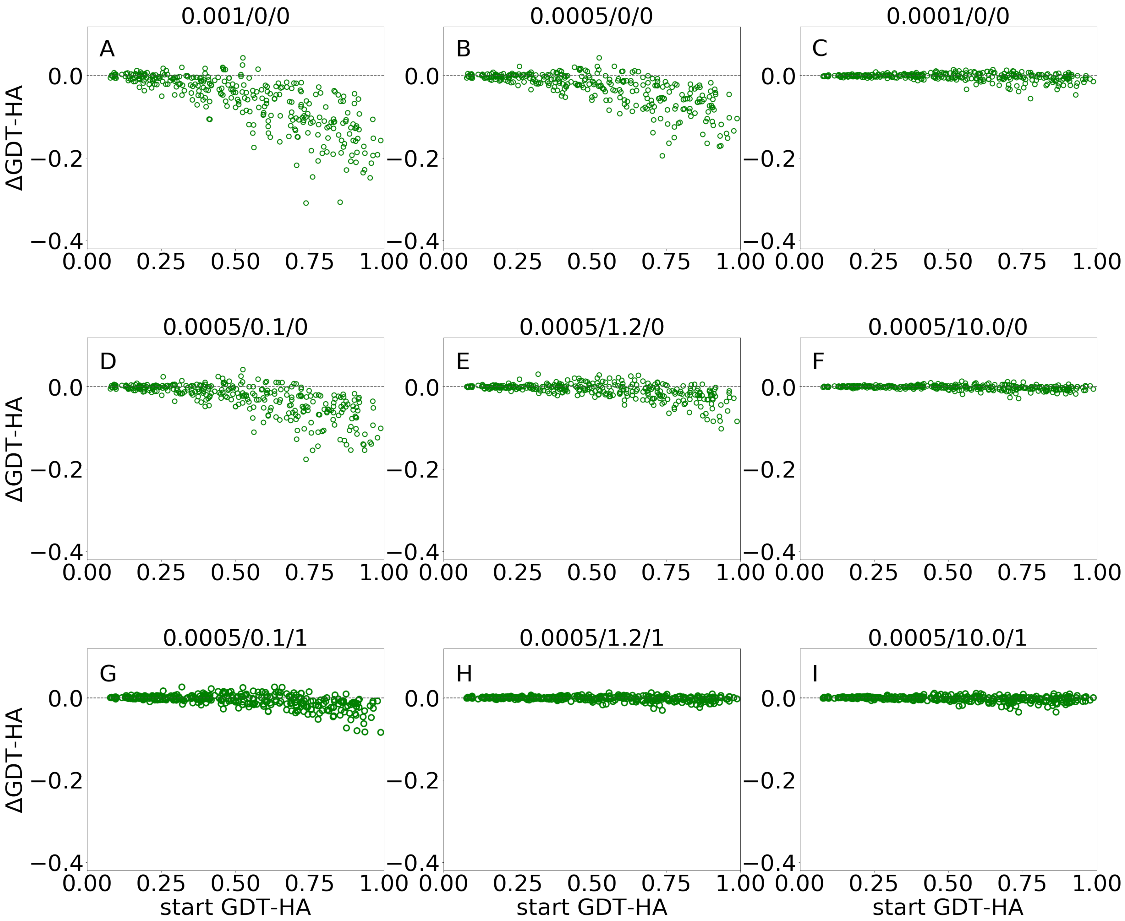


Figure S5: Scatter plots of ΔGDT-HA as a function of start GDT-HA for top 1 models by GSFE-Refinement for various /LR/$\lambda$/W combinations on 3DRobot dataset

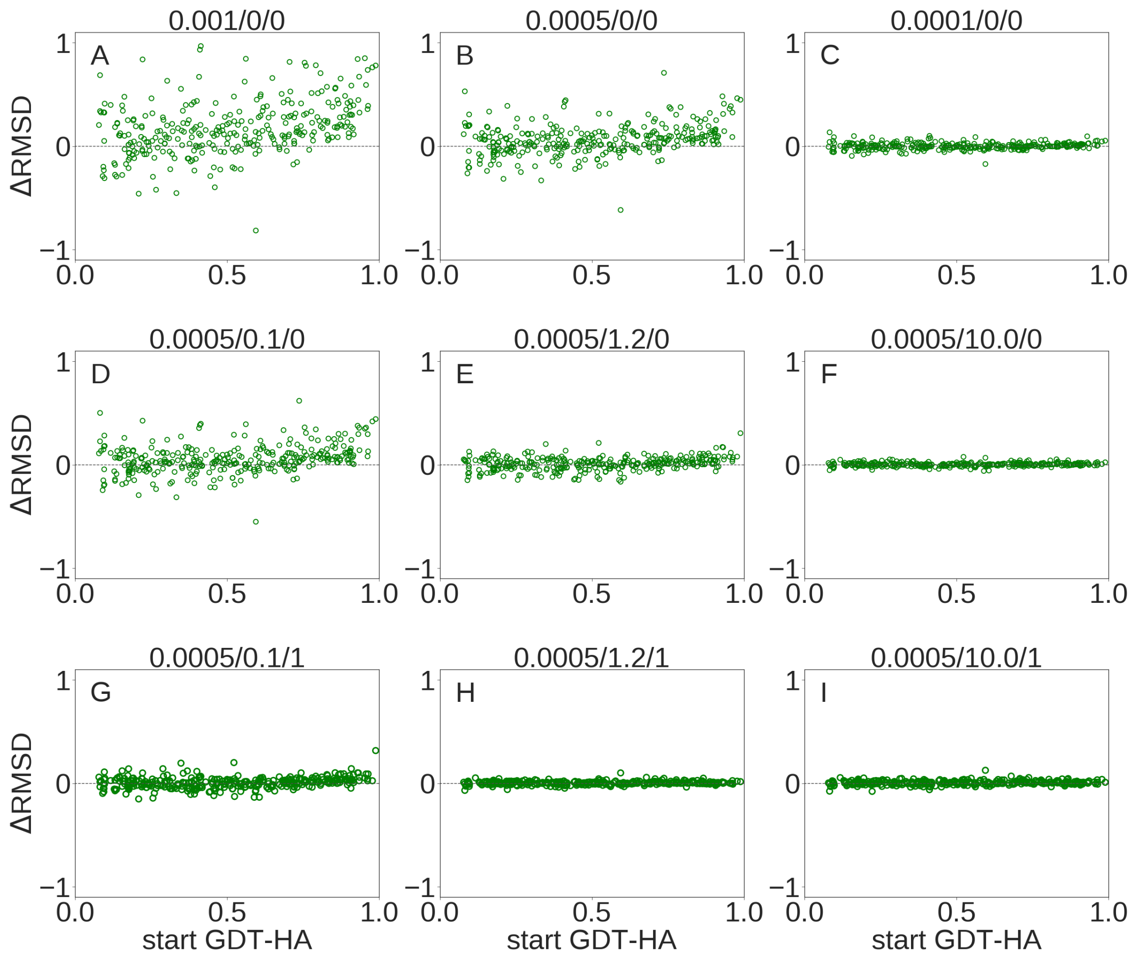


Figure S6: Scatter plots of ΔRMSD as a function of start GDT-HA for top 1 models by GSFE-Refinement for various /LR/$\lambda$/W combinations on 3DRobot dataset

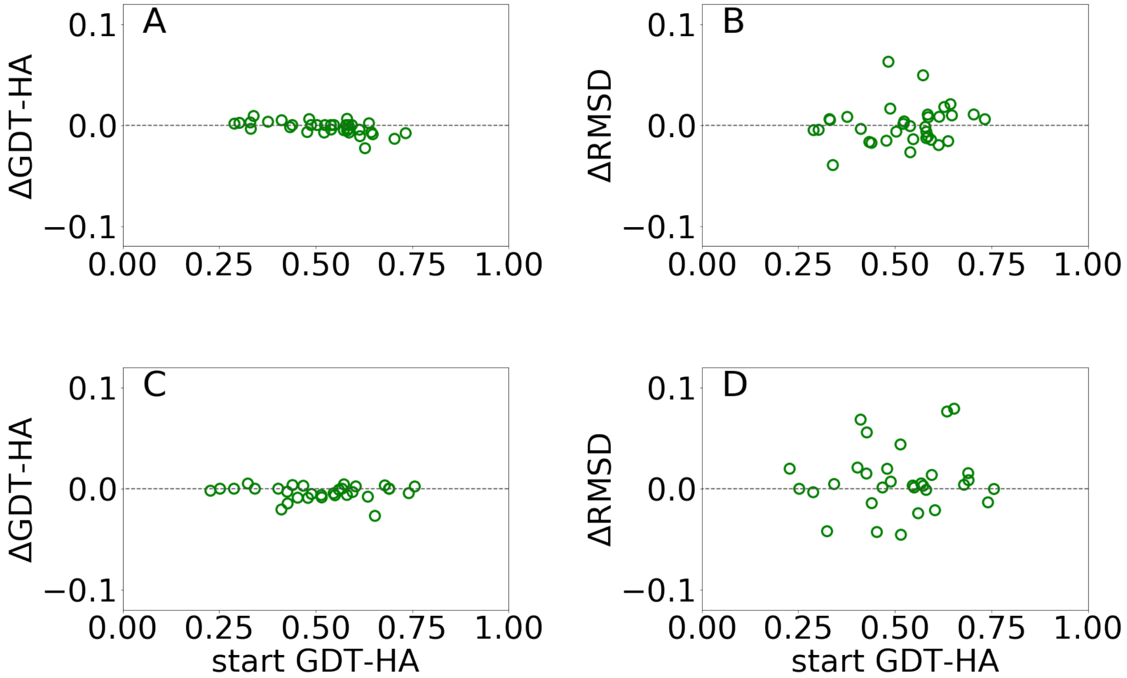


Figure S7: Scatter plots of ΔGDT-HA (A,C), ΔRMSD (B,D) as a function of start GDT-HA for top 1 models by GSFE-Refinement for various /LR/λ/W combinations on CASP11 and CASP12 dataset

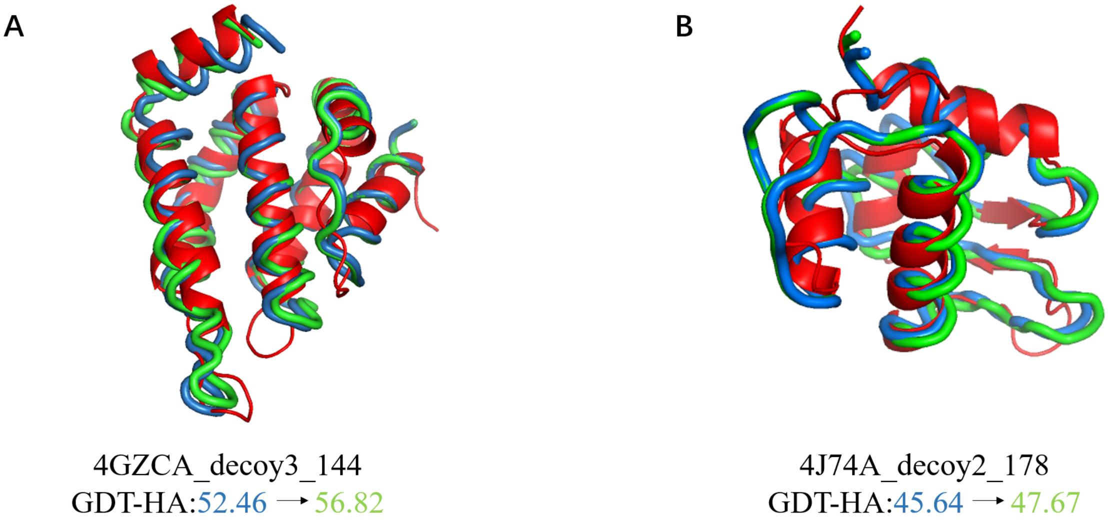


Figure S8: Two examples of GSFE-refinement with larger improvement for learning rate 0.0005 than 0.0001. The starting (red) and refined (green) decoys are overlaid on the experimental (blue) structures with change of GDT-HA at learning rate of 0.0005 indicated. Corresponding change of GDT-HA is 0.57 for4GZCA_decoy3_144 and 0.58 for 4J74A_decoy2_178 respectively.

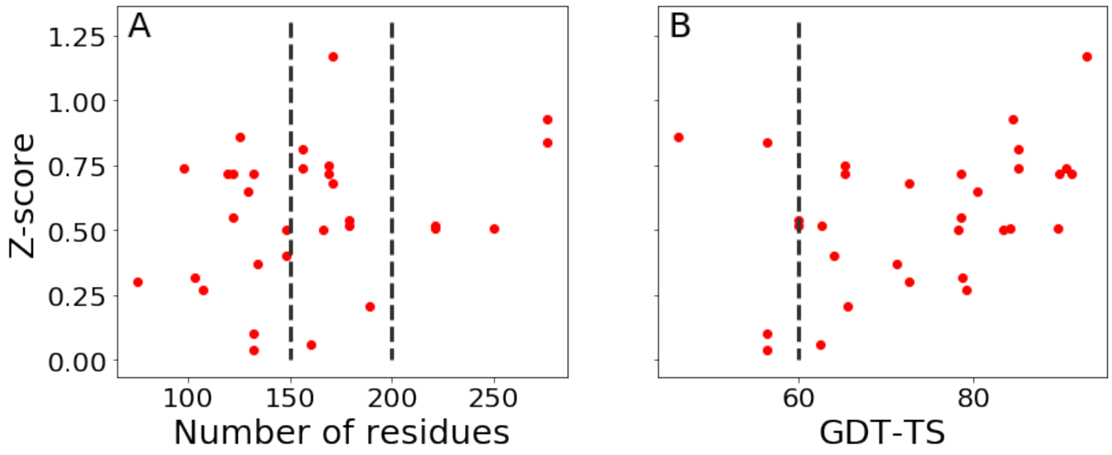


Figure S9: GDT-TS based Z-scores for top 1 model by GSFE-Refinement in CASP14. (A) As a function of the number of residues. (B) As a function of starting GDT-TS score.
